# Computational modeling predicts acidic microdomains in the glutamatergic synaptic cleft

**DOI:** 10.1101/2021.06.30.450624

**Authors:** Touhid Feghhi, Roberto X. Hernandez, Michal Stawarski, Connon I. Thomas, Naomi Kamasawa, A.W.C. Lau, Gregory T. Macleod

## Abstract

At chemical synapses, synaptic vesicles release their acidic contents into the cleft leading to the expectation that the cleft should acidify. However, fluorescent pH probes targeted to the cleft of conventional glutamatergic synapses in both fruit flies and mice reveal cleft alkalinization, rather than acidification. Here, using a reaction-diffusion scheme, we modeled pH dynamics at the *Drosophila* neuromuscular junction (NMJ) as glutamate, adenosine triphosphate (ATP) and protons (H^+^) are released into the cleft. The model incorporates bicarbonate and phosphate buffering systems as well as plasma membrane calcium-ATPase (PMCA) activity and predicts substantial cleft acidification but only for fractions of a millisecond following neurotransmitter release. Thereafter, the cleft rapidly alkalinizes and remains alkaline for over 100 milliseconds, as the PMCA removes H^+^ from the cleft in exchange for calcium ions (Ca^2+^) from adjacent pre- and post-synaptic compartments; thus recapitulating the empirical data. The extent of synaptic vesicle loading and time course of exocytosis has little influence on the magnitude of acidification. Phosphate, but not bicarbonate buffering is effective at ameliorating the magnitude and time course of the acid spike, while both buffering systems are effective at ameliorating cleft alkalinization. The small volume of the cleft levies a powerful influence on the magnitude of alkalinization and its time course. Structural features that open the cleft to adjacent spaces appear to be essential for alleviating the extent of pH disturbances accompanying neurotransmission.

**SIGNIFICANCE STATEMENT:** Acid-base imbalances have surprisingly potent neurological effects highlighting the acute pH sensitivity of many neural mechanisms. Acid-Sensing Ion-Channels (ASICs), which open in response to acid shifts in extracellular pH, are an example of such a mechanism. However, while ASICs open during neurotransmission at conventional glutamatergic synapses, pH-sensitive electrodes and fluorophores show no signs of acidification at these synapses, only alkalinization. To resolve this paradox, we built a computational model which allows a glimpse beyond the experimental limitations of pH-sensitive electrodes and fluorophores. Our model reveals a highly dynamic pH landscape within the synaptic cleft, harboring deep but exceedingly rapid acid transients that give way to a prolonged period of alkalinization, thus reconciling ASIC activation with direct measurements of extracellular pH.

## INTRODUCTION

Any doubt that our nervous system is pH sensitive is put to rest by the neurological consequences of even the smallest systemic acid/base disturbance [1–5]. The degree to which nerve activity itself can drive pH change is poorly understood yet any such change would likely impact the efficacy of neurotransmission as voltage-gated calcium channels (VGCCs), glutamate receptors (GluRs), gamma-aminobutyric acid type-A receptors (GABA_A_R) and acid-sensing ion channels (ASICs) are exposed at the cleft and are pH-sensitive at their extracellular aspect [1–3, 6].

The preponderance of electrophysiological data indicate acidification at vertebrate sensory ribbon synapses during neurotransmission [7–11]. Fluorescent pH reporters targeted to the cleft at these synapses also reveal acidification [12, 13]. However, conventional neuronal synapses appear to be different, as fluorescent pH reporters targeted to such synapses in mice and fruit flies reveal cleft alkalinization without any signs of acidification [14]. Furthermore, alkalinization has also been observed adjacent to glutamatergic synapses in the mammalian hippocampus and cerebellar cortex [15–22]. Yet, it is clear that ASICs are activated at conventional glutamatergic synapses within the amygdala, nucleus accumbens, calyx of Held and cingulate cortex [23–26]. Clearly, these reports of ASIC activation are difficult to reconcile with pH recordings made beside these synapses and within their clefts.

Here we adopted a computational approach to model competing influences on pH within the cleft at the *Drosophila* NMJ. The chemical synapse, the fundamental unit of intercellular communication and neural computation, represents a challenge for any pH buffering system. It is characterized by a site of close apposition between pre- and post-synaptic neurons; the synaptic cleft. Close apposition is necessary to deliver a package of neurotransmitters at high concentration onto corresponding receptors on the surface of the post-synaptic neuron. However, this confined space is expected to manifest substantial acidification caused by the release of the conjugate acid of neurotransmitters within synaptic vesicles (SVs), and the limited pH buffering entities present in small interstitial volumes. In opposition to this acidifying influence is the PMCA, a Ca^2+^/2H^+^ exchanger, which exerts an alkalinizing influence on interstitial spaces [27]. Ca^2+^ enters the presynaptic compartment through VGCCs and Ca^2+^ enters the postsynaptic compartment through neurotransmitter-gated channels, while the PMCA removes Ca^2+^ from both compartments in exchange for H^+^ from interstitial spaces.

Using a reaction-diffusion scheme implemented in MATLAB, we modeled the effects on cleft pH of neurotransmitters and their conjugate acids as they are released from SVs, interact with buffers, and diffuse away from the release site. Furthermore, we incorporated into this model the effect of PMCA and the effect of opening the synaptic cleft to the voids within the adjacent postsynaptic folds. We found that this conventional glutamatergic synapse will, as expected, acidify. However, acidification only lasts for fractions of a millisecond followed by a predominant alkalinization that summates during burst firing. In addition, we observed that opening the cleft to adjacent spaces greatly alleviates the extent of pH disturbances accompanying neurotransmission.

## RESULTS

### Imaging data show that the *Drosophila* NMJ synaptic cleft alkalinizes during activity

The NMJ of *Drosophila* larvae is characterized by axonal swellings (boutons) making multiple synapses with a highly elaborated postsynaptic membrane [28]. Our ability to target pH-sensitive fluorescent proteins to the outside surface of either pre- or postsynaptic membranes has enabled us to monitor pH changes within the synaptic cleft (**Fig. 1A-B**) [14, 29]. Without exception, the change in pH in response to a single AP (ΔpH_AP_) is a rapid alkalinizing transient (**Fig. 1C**). The average ΔpH_AP_ is ~0.025 log units, peaking ~50 ms after the nerve impulse, and decaying with a time course of ~140 ms [14]. The suppression of all signs of pH change upon the addition of 7 mM glutamic acid [**Fig. 1C**; which causes postsynaptic glutamate receptor desensitization [30]], or when Ca^2+^ is omitted from the bath saline (data not shown), led us to conclude that Ca^2+^ entry to the postsynaptic muscle and its subsequent extrusion by PMCA drives cleft alkalinization [14]. The change in pH in response to a train of stimuli (ΔpH_train_), intended to simulate the axon’s activity during a peristaltic body-wall contraction, was well in excess of ΔpH_AP_ (**Fig. 1D**) and was similarly occluded by the presence of glutamate or absence of Ca^2+^ (data not shown). Our broad interpretation of the ion fluxes that drive a single AP evoked alkaline transient is sketched in **Figure 1E**.

**Figure 1.**
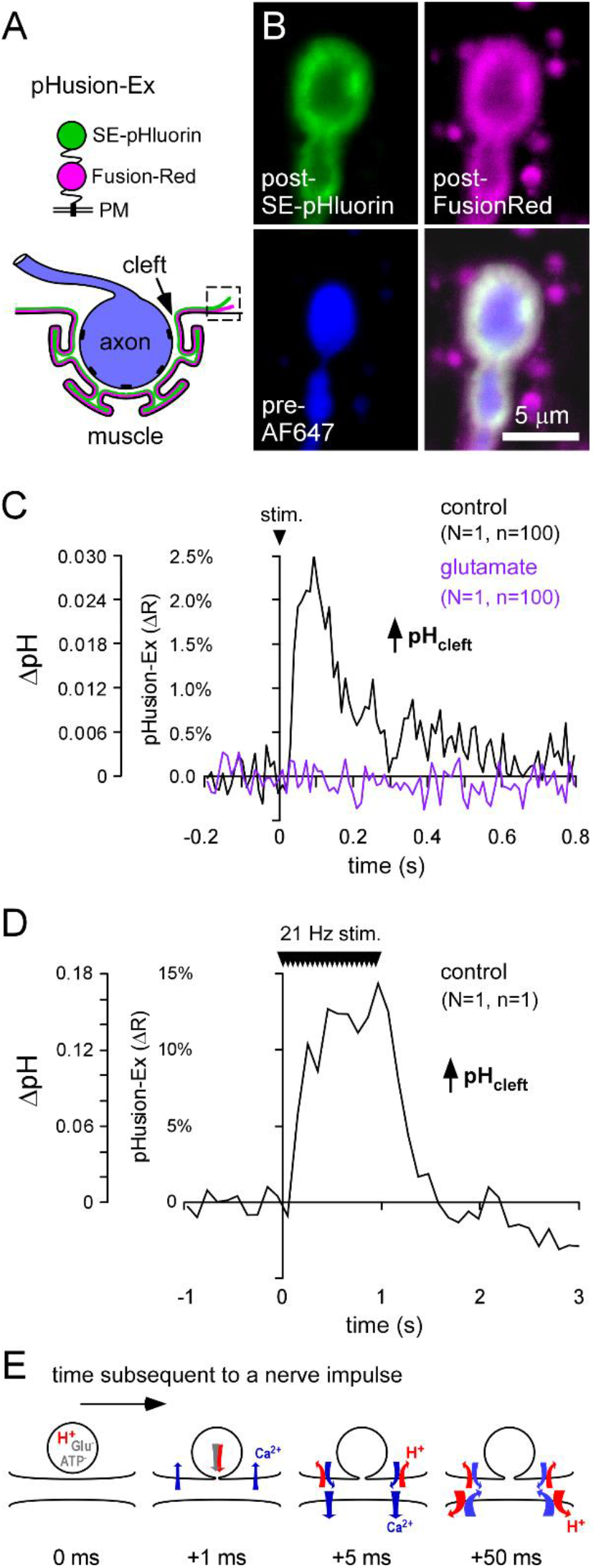
The cleft of the *Drosophila* NMJ alkalinizes during activity. (**A**) The pH sensitive pHusion-Ex probe can be targeted to the extracellular spaces of the NMJ as an anchor using a transmembrane domain. (**B**) When expressed in the muscle the probe is seen in the SSR that surrounds motor nerve terminal boutons (revealed here through forward-filling the live axon with AF647). (**C**) Cleft alkalinization can be seen in response to a single AP, providing signal averaging is used. Both traces represent the average of 100 APs delivered at 1 Hz. (**D**) Cleft alkalinization can be seen in response to a train of APs (without signal averaging) delivered at the neuron’s native firing frequency (21 Hz). In C and D, NaHCO_3_ is added to 15 mM and continuously bubbled with carbogen; stabilizing at pH 7.2. No phosphate or zwitterionic buffers present. (**E**) Our interpretation of this phenomenon is summarized in a temporal series: In response to an AP, Ca^2+^ entry to the pre-synaptic terminal triggers the exocytosis of glutamate, co-loaded ATP, and protons titrated by both. Activation of post-synaptic glutamate receptors allows Ca^2+^ entry to the pre-synaptic muscle. The PMCA extrudes Ca^2+^ from both pre- and post-synaptic compartments using protons as a counter-ion.

### The NMJ is characterized by a narrow cleft and high PMCA activity

In an attempt to reconcile our empirical data with both the release of H^+^ from SVs and reports of ASIC activation at conventional glutamatergic synapses, we proceeded to a theoretical treatment of pH dynamics inspired by a previous numerical modeling approach [31]. As a first step, we measured critical distances at the synapse to establish geometrical constraints. Synaptic boutons form multiple AZs, readily visualized through immunohistochemistry (**Fig. 2A**).

**Figure 2.**
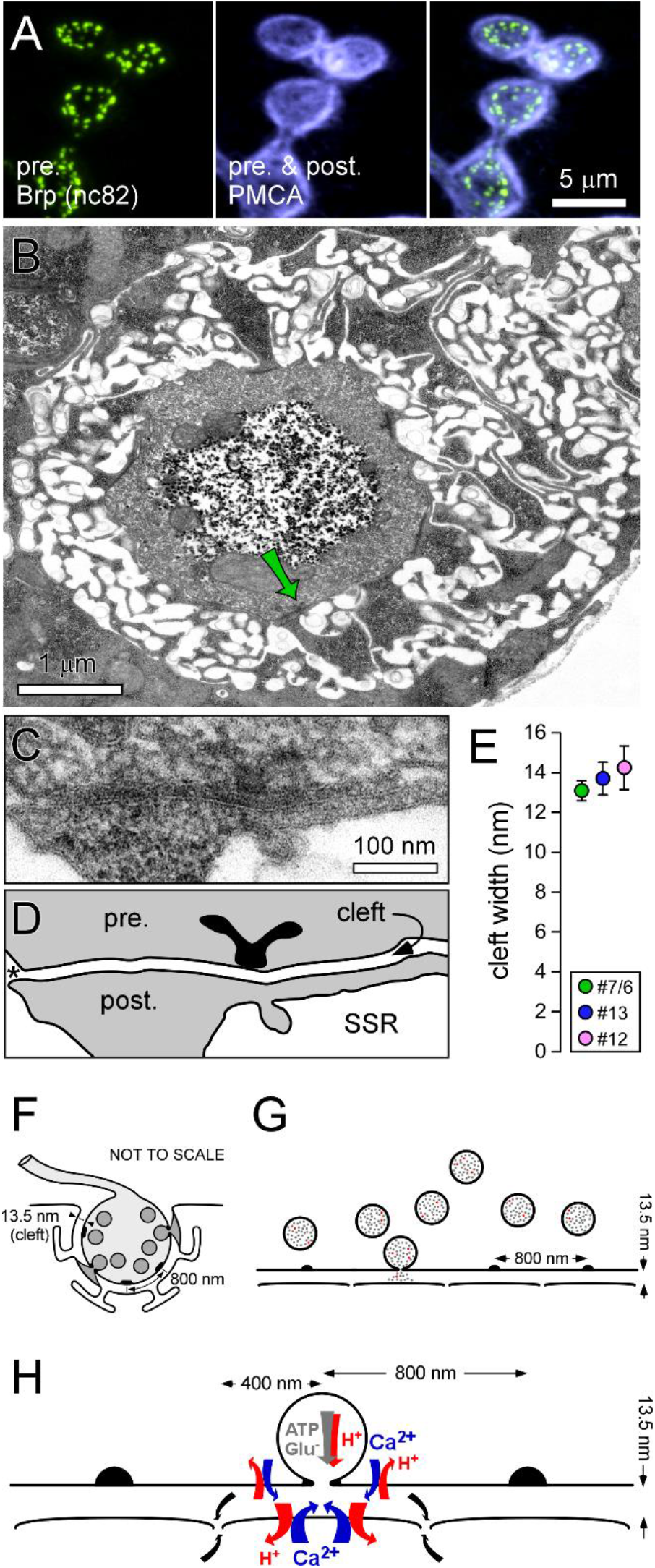
The *Drosophila* NMJ synaptic cleft is very narrow, but continuous with an adjacent void. (**A**) Fluorescent images of two proteins at the NMJ revealed by immunohistochemistry; Presynaptic Bruchpilot (Brp), defining the AZs, and PMCA found in both pre- and postsynaptic compartments. (**B**) An electron micrograph of a 50 nm section through the center of a presynaptic bouton and the surrounding muscle SSR. AZs, which are often marked by a “T-bar” extending into the cytosol from the pre-synaptic membrane (green arrow), can be found at a number of sites around the bouton’s periphery. (**C**) An enlarged view of the AZ indicated by the green arrow in B. (**D**) Interpretation of the image in (**C**), emphasizing the location of the T-bar, the cleft, and the common observation that the cleft usually empties out into voids of the SSR ~ 400 nm distant from the center of each AZ (see asterisk). (**E**) Plot of the width of the cleft (between outer leaflets of opposing plasma membranes) at NMJs on muscle fibers #7/6, 13 and 12. Each point represents measurements at 12 AZs at each of 5 boutons on each of the muscle fibers; mean ± SEM. (**F)** Schematic of a cross-section of a bouton embedded within the muscle SSR, showing release into the cleft from 2 out of 5 AZs. (**G**) For computational purposes, the cleft can be treated as a continuous layer, sandwiched between the planes of two opposing membranes; represented here in cross section annotated with critical distances. (**H**) An expanded view of release and subsequent PMCA activity, emphasizing the location of PMCA activity and the location at which the cleft becomes continuous with the voids within the SSR.

PMCA is heavily concentrated in the postsynaptic folds, and, while its presynaptic localization cannot be distinguished from the postsynaptic immunofluorescence signal, Ca^2+^ imaging studies have demonstrated the critical role of PMCA in extruding presynaptic Ca^2+^ in addition to postsynaptic Ca^2+^ [32, 33]. AZ spacing has been quantified in numerous EM studies indicating an average spacing of ~0.8 μm in 3^rd^ instar larvae [34–36]. In contrast, there are few reports of cleft width, thus we examined thin sections through larval 3^rd^ instar NMJs on muscle fibers #6/7, 13 and 12. We defined cleft width as the distance between the outer-leaflet of the postsynaptic compartment and the outer-leaflet of the presynaptic compartment, yielding an average width of 13.5 ± 0.4 nm (**Fig. 2B-E**). These values are generally consistent with other measurements made at the 3^rd^ instar NMJ [17 ± 1 nm; [37]), with measurements via EM tomography in 2^nd^ instar larvae prepared through high-pressure freeze freeze-substitution [15.9 ± 0.1 nm; [38]], and with reconstructions of super-resolved synaptic proteins [39]. Our EM examination also revealed that close apposition of the postsynaptic membrane often extends no further than ~0.4 μm from either side of the center of each AZ (arrow) before opening out into the interstitial space of the SSR (**Fig. 2D**). This can also be observed in previously published images of chemically fixed NMJs [36, 40].

### An overview of the computational model

The geometry of our model is shown in **Figure 2F-H**, along with the identity of some of the molecular species and their relative movement. Briefly, we approximate the cleft as a gap between two infinite planes, each plane representing the outer face of the pre- and postsynaptic compartment. We define the center of the AZ as the site of exocytosis. Once the acidic contents of a SV are released into the cleft, both Glu and ATP titrated with H^+^, diffuse throughout the cleft, becoming more dilute while releasing H^+^ that react with bicarbonate and phosphate buffers. At pre- and postsynaptic compartments, the PMCA mediates the exchange of H^+^ in the cleft for Ca^2+^ in pre- and postsynaptic compartments. Using coupled reaction-diffusion equations, as shown in **Equation 1**, we calculate the spatio-temporal concentration of each of 12 species within the cleft. See **Table 1** for the identity of all species. We denote the concentration of a particular species as function of the location, r, and time, t by c_i_(r, t), where i = 1 to 12. For example, the concentration of H^+^ is denoted as [H^+^] = c_1_(r, t).

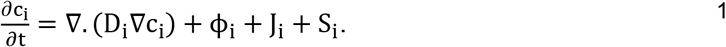

**Table 1.** Identity of all molecular species accommodated by Eq. 1.

|  |  |  |
| --- | --- | --- |
| $[H^+] = c_1(r, t)$ | $[HCO_3^-] = c_2(r, t)$ | $[H_2CO_3] = c_3(r, t)$ |
| $[CO_2] = c_4(r, t)$ | $[H_2PO_4^-] = c_5(r, t)$ | $[HPO_4^{2-}] = c_6(r, t)$ |
| $[H_2Glu] = c_7(r, t)$ | $[HGlu^-] = c_8(r, t)$ | $[Glu^{2-}] = c_9(r, t)$ |
| $[H_2ATP^{2-}] = c_{10}(r, t)$ | $[HATP^{3-}] = c_{11}(r, t)$ | $[ATP^{4-}] = c_{12}(r, t)$ |

Each term on the right-hand side of **Equation 1** is discussed, along with assumptions, in the Methods. Briefly, the first term *D_i_* captures Fick’s law of diffusion for each species; the diffusion coefficient for species i. The second term ϕ_i_ is the reaction term of species i, representing the production and consumption of that species as given in **Equations 9–20** in the Methods. The third term, *J_i_* denotes the extrusion of H^+^ from the cleft through PMCA which exchanges Ca^2+^ with H^+^ while maintaining electroneutrality, i.e., 2H^+^ for Ca^2+^. The last term *S_i_* represents the source term of each species.

### Cleft acidification is substantial, but fleeting

The model was first used to explore cleft pH at the site of exocytosis where, under nearly all conditions, it revealed a deep but short-lasting acid transient (red trace, **Fig. 3A**). With bicarbonate present at 19 mM, facilitated by medium CA activity (CA; 600 nominal units), cleft pH plunged to 5.5 within microseconds of exocytosis. Perhaps the most surprising result was the ephemeral nature of cleft acidification. Even without bicarbonate or phosphate in the cleft, acidic transients “evaporated” within 500 μs through diffusion alone (**Fig. 3B**). When both buffering systems were enabled, cleft acidification was extinguished within only 50 μs. Equidistant between the site at which exocytosis occurs and the site at which the cleft gives way to the SSR, bicarbonate buffering greatly ameliorated the depth of the acidic transients and total phosphate buffering of 1 mM almost extinguished them (**Fig. 3C**). At the site of release, the cleft invariantly acidified to the same pH as the SV lumen, independent of buffering conditions. Similarly, assumptions about SV loading in a range of 1,500 to 8,000 glutamate molecules, with or without co-loaded ATP at 100 mM (610 molecules), were not influential with regard to the depth of acidification at the site of release (**Fig. 3D**). The time course of exocytosis has been estimated to be less than 100 μs [41, 42] but changing the time course of Glu release in this realm (0.1 − 100 μs) has little influence on the depth of acidification or the rate at which the cleft recovers to neutral (**Fig. 3E&F**). These data indicate that cleft acidification is inevitable at the site of exocytosis, and that it is of sufficient magnitude and duration to activate ASICs [43, 44]. Understandably, signs of cleft acidification terminating within fractions of a millisecond will be missed by imaging systems sampling at rates as slow as ~ 10 ms per frame (100 Hz, **Fig. 1C**) or 2 ms per line (500 Hz, [14]).

**Figure 3.**
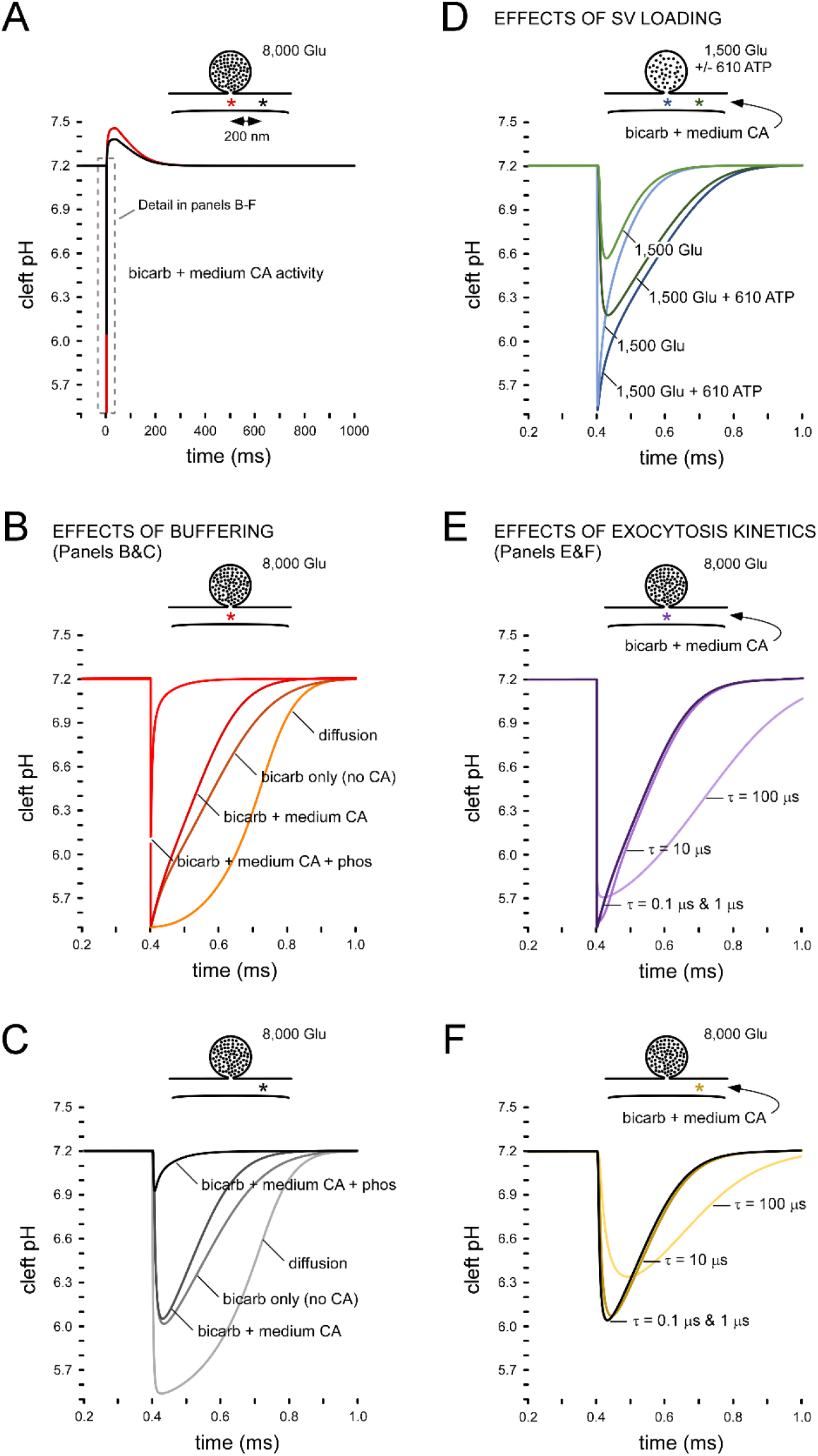
The synaptic cleft shows substantial but brief acidification. (**A**) Output plots of the computational model. When 8,000 glutamate molecules are released into the cleft containing 19 mM bicarbonate and a medium level of CA activity [enzymatic acceleration (En_acc_ of 600 (nominal units)], the pH at the mouth of fusion pore drops to 5.5 (i.e. equilibrates with SV contents) within microseconds (deep red trace – corresponding to location of the deep red asterisk). The cleft then alkalinizes, reaching a peak of ~ 7.45 within ~50 ms, before decaying to the pre-stimulus baseline at pH 7.2. Midway between the fusion pore and access to the SSR (location of the black asterisk), both acidification and alkalinization are ameliorated in magnitude (black trace). (**B-C**) Bicarbonate buffering has little impact on cleft acidification at either the site of release or midway between the fusion pore and SSR; phosphate buffering, however, is very effective (1 mM total). (**D**) If SV loading is reduced to 1,500 glutamate so that it is isosmotic with the cytosol (250 mM), and co-loaded with 610 ATP molecules (100 mM; total 350 mM), the acid transient is similarly deep as when 8,000 glutamate is released. (**E-F**) The time course of fusion pore opening similarly has little impact on the depth of acidification over a range of 4 orders of magnitude (τ = 0.1−100 μs; 8,000 glutamate). Bicarbonate buffering is 19 mM in all scenarios except diffusion only. Fusion pore opening (τ) is 1 μs, except where indicated otherwise (E&F).

### Ca^2+^ pumping across membranes alkalinizes the cleft

An exploration of cleft pH for time periods beyond SV exocytosis required that PMCA activity be incorporated into the model and this revealed cleft alkalinization consistent with our empirical data (**Fig. 4A-D**). With bicarbonate present at 19 mM, and a “medium” level of CA activity [similar to conditions used in **Fig, 1C&D**, and throughout Stawarski et al., (2020)], PMCA mediated exchange of cleft H^+^ for cytosolic Ca^2+^ resulted in cleft alkalinization up to 7.45, from a baseline of 7.20 (ΔpH ~0.25), peaking ~50 ms after exocytosis of a single SV. Cleft alkalinization begins at a very low rate once PMCA commences on the presynaptic plasma membrane and this can be seen in **Figure 4D**, right panel (#), prior to exocytosis (†). Alkalinization accelerates as the activity of postsynaptic PMCA comes online [Fig. 4B&C (right panels; arrows) and Fig. 4D (ψ))]. The extent of alkalinization was sensitive to the degree of CA facilitation and almost extinguished by the presence of 1 mM total phosphate. A ΔpH of 0.25 is much greater than estimates made from empirical data (0.02; Fig.1C; [14]), but empirical estimates rely on a signal-averaging technique that occludes estimates of ΔpH at individual AZs (Illustrated in **Fig. 6**). We and others have previously established that the average probability of release from the AZs along these terminals is ~0.11 [45, 46]. Therefore, as the empirical estimate of 0.025 was generated from groups of AZs where exocytosis occurs with a probability of only ~0.11, ΔpH for a single AZ undergoing exocytosis might be recalculated as 0.23 (1/0.11 × 0.025), consistent with the displacement predicted by the model (~ 0.25). While cleft alkalinization does not reach magnitudes as great as acidification during exocytosis, the model shows that it persists for several orders of magnitude longer.

**Figure 4.**
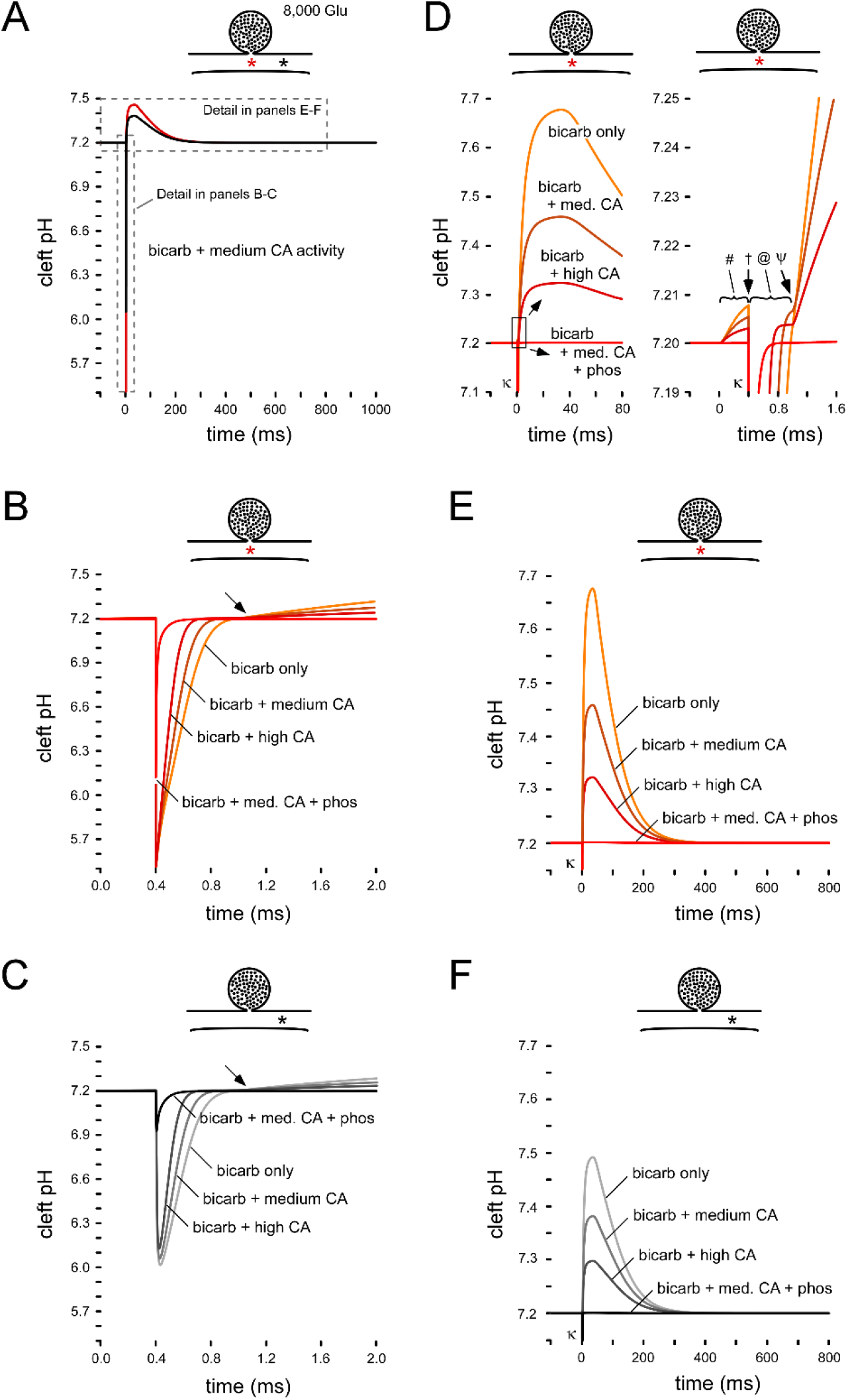
The synaptic cleft shows moderate but long lived alkalinization. (**A**) Identical to figure 3A, the model output plots in panel A provide context for the detail of B-F. (**B-C**) show the minimal impact of bicarbonate buffering on cleft acidification caused by neurotransmitter release. The extended time-scale encompasses the transition point from acidification to alkalinization (ψ) at ~0.9 ms (when the impact of post-synaptic PMCA activity becomes apparent). (**D**) Left panel shows the first 80 ms after release, illustrating the shape of alkaline transient which does not decay until after post-synaptic Ca^2+^ entry ceases at ~40 ms. Right panel shows detail of the boxed section on the left. # : cleft alkalinization as a result of pre-synaptic PMCA activity; † : cleft acidification as a result of exocytosis; @ : realkinization as a result of diffusion, buffering and pre-synaptic PMCA activity; ψ : alkalinization as a result of pre- and post-synaptic PMCA activity. (**E-F**) Bicarbonate, and especially phosphate buffering (1 mM), is effective at ameliorating cleft alkalinization. Note: the 400 fold change in time base between B-C and E-F. 8,000 glutamate molecules released and 19 mM bicarbonate buffering in all scenarios. Fusion pore opening (τ) is 1 μs. Medium and high CA activity are 600 and 2,000 En_acc_ nominal units, respectively. κ indicates truncated acidic transients.

### Synapse morphology has a powerful influence on cleft pH

While motor neuron axon terminals are buried in the muscle at the *Drosophila* NMJ, the cleft never-the-less opens out into voids within the post-synaptic folds, i.e. within the SSR. This space represents a large extracellular volume in close contact with a large area of muscle plasma membrane. Here we used the model to explore the role of the SSR in cleft pH homeostasis. The 12 ionic species considered in this model are restricted to move and react within the cleft, but for several exceptions: CO_2_ is allowed to diffuse freely across membranes, while Ca^2+^ and H^+^ can move across membranes via VGCCs and GluRs (Ca^2+^) and PMCA (Ca^2+^ in exchange for 2H^+^). The consequences of not allowing species to exchange with the SSR 0.4 μm distant from the release site is shown in **Fig. 5**. Providing CA is present to support the buffering provided by 19 mM bicarbonate, pH can recover to neutral after SV exocytosis. However, below a certain threshold of CA activity (~500 nominal units), pH becomes unstable as the PMCA extrudes H^+^ to an extent where PMCA will begin to stall only milliseconds after activation (pH ~8.8) (**Fig. 5A**). Distance from the site of release has little consequence (**Fig. 5B**). Time to stall is greatly abbreviated if the free movement of CO_2_ is prevented (**Fig. 5C-D**), as cleft pH becomes unstable in tens of microseconds, an outcome that can be slowed but not prevented by phosphate buffering. Blockage of species exchange with the SSR, or the free movement of CO_2_, has little impact on the magnitude of cleft acidification, and only a moderate effect on the duration of acidification (**Fig. S4**).

**Figure 5.**
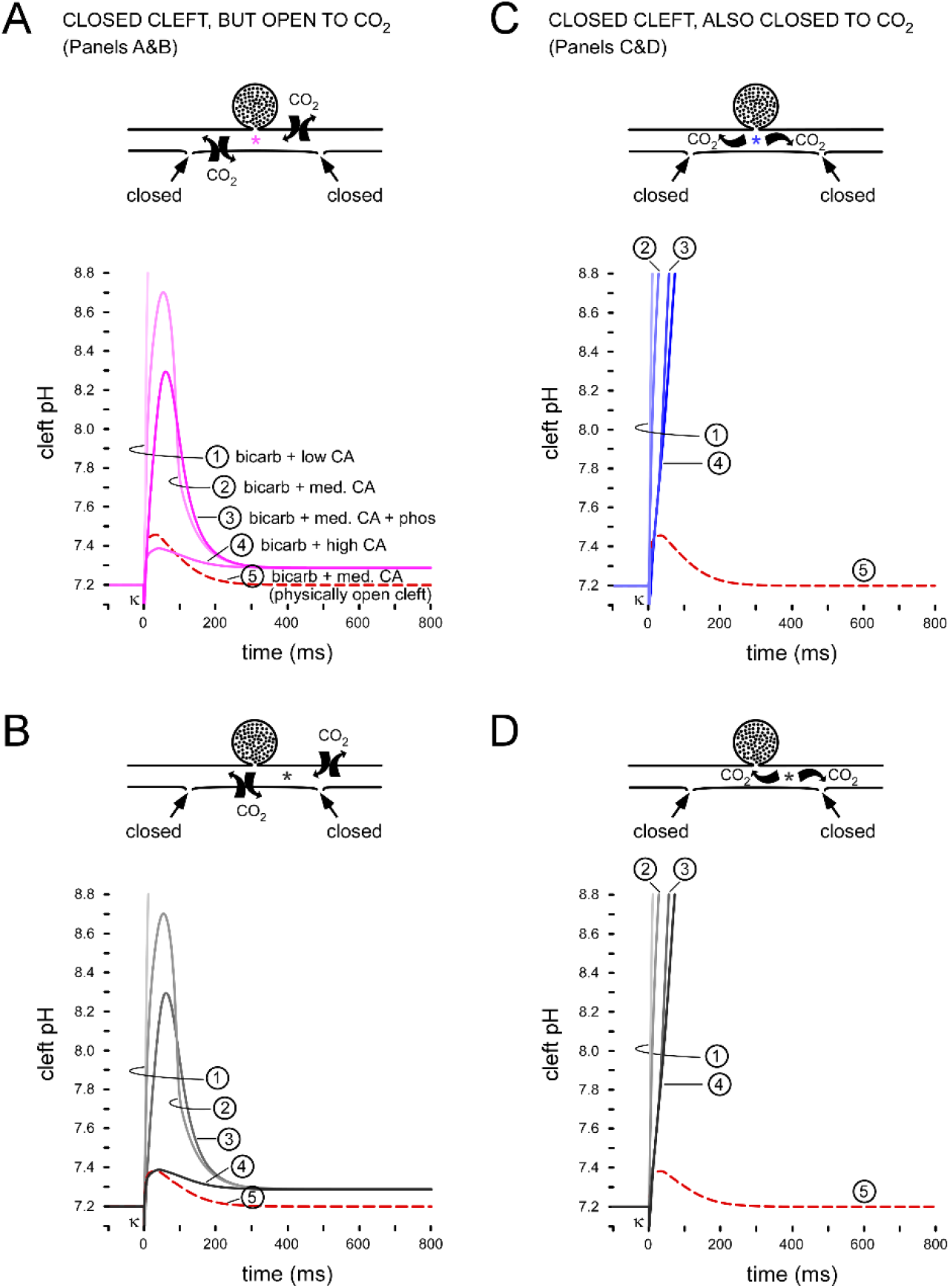
Synapse morphology has a powerful influence on cleft pH. (**A-B**) Output plots of the computational model showing the impact of preventing access from the synaptic cleft to the void of the SSR (note the location of asterisks in A vs B). The dashed line represents the control conditions, i.e. bicarbonate buffering in the presence of medium CA activity, full access to the SSR and membrane permeability to CO_2_. Low, medium and high CA activity are 100, 600 and 2,000 En_acc_ nominal units, respectively. Note, because of the ability of CO_2_ to cross membranes, high CA activity is better able to buffer pH changes than medium CA activity in combination with 1 mM total phosphate. When CA activity (En_acc_) falls below ~500 nominal units cleft buffering fails. pH 8.8 represents the pH at which the PMCA is expected to stall [33]. (**C-D**) Plots showing the impact of both preventing cleft access to the SSR, and preventing CO_2_ movement across membranes. Cleft pH buffering failed in all cases (note the location of asterisks in C vs D). 8,000 glutamate molecules released and 19 mM bicarbonate buffering in all scenarios. Fusion pore opening (τ) is 1 μs. κ indicates truncated acidic transients. Detail of acidic transients and first 80 ms of the alkaline transients are given in Figure S4.

### Cleft alkalinization will summate at high frequencies

As the cleft acidifies for such a short period (<1 ms) it is highly unlikely that the axon would fire at a rate rapid enough (>1 kHz) for vestiges of acidification to persist in the cleft until the arrival of a subsequent AP. Alkalinization, however, is relatively prolonged, in which case vestiges of alkalinization might persist between APs. The model was used to examine the axon firing frequencies at which alkalinization begins to summate at an AZ (**Fig. 6**). Contrary to the expectation given by the buildup of alkalinization in **Figure 1D**, the model showed no signs of pH summation at individual AZs until axon firing well exceeded its native frequency of 21 Hz. This counterintuitive result is a function of pH change being insulated between AZs and the probabilistic nature of neurotransmitter release. Release occurs with an average probability of only ~0.11 at these AZs [45, 46], in which case, although APs arrive in the presynaptic terminal at 21 Hz, release events occur at an average rate of 2.3 Hz (21 × 0.11) at any particular AZ. Insufficient alkalinization persists in the cleft when release occurs at only 2.3 Hz, i.e. with an interval of 435 ms between release events. At axon firing frequencies beyond ~ 40 Hz alkalinization summates at individual AZs (**Fig. 6D-E**). At 40 Hz, the “average” AZ would release at ~4.4 Hz (40 × 0.11), i.e. once every ~ 230 ms, and at this rate 7% of the alkaline transient persists from the previous event. To the extent that fictive firing rates represent axon firing rates during locomotion, these axons fire at a rate below the threshold required to obtain a “boost” from a gain mechanism that exploits cleft alkalinization.

**Figure 6.**
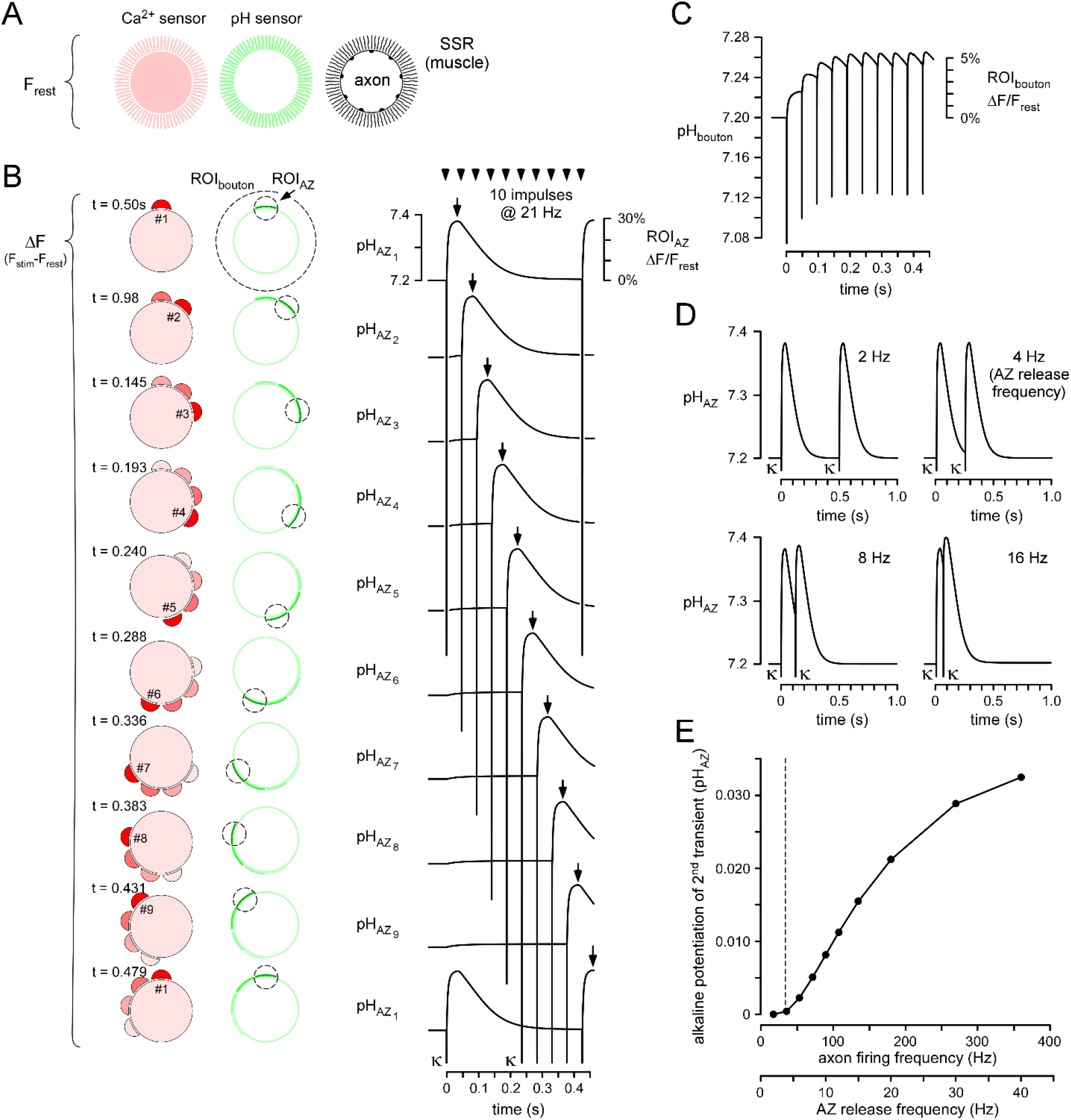
MNs must fire beyond usual rates before alkaline transients will summate at individual AZs. (**A-C**) A scheme which illustrates why pH reporter fluorescence summation from a bouton over multiple nerve impulses may not indicate pH transient summation at individual AZs. (**A**) Stylized representation of resting fluorescence of a cytosolic Ca^2+^ reporter (both sides of the synapse) and an extracellular pH reporter (muscle PM) at a single bouton of the NMJ. (**B**) Representation of the Ca^2+^ and pH responses to release from individual AZs over 10 APs; Ca^2+^ response are postsynaptic and pH responses are in the cleft. The Ca^2+^ reporter responses are shown to delineate the location of each releasing AZ. The average probability of release is 1/9, and 9 AZs have been illustrated responding sequentially around the periphery of a bouton. The images capture the maximal response; the time of image capture is represented with an arrow on the pH plots for each AZ. The plots of pH change at each of the AZs are generated by the model (LHS ordinate). The images show the location of an AZ-delimited region-of-interest (ROI) that would be expected to yield pH sensor transients (ΔF/F_rest_) of the magnitude shown (RHS ordinate). **C**, Numerical summation of all AZ pH plots in B yields a pH change for the entire bouton that is well below that of an AZ, but, unlike pH at an individual AZ, it summates over multiple APs. The RHS ordinate shows the expected fluorophore transients (ΔF/F_rest_) if the ROI encompassed the entire bouton containing the 9 AZs, and this is compatible with what is observed empirically (Fig. 1D). (**D**) Model generated plots of pH an individual AZ where one release event follows another after a defined interval. (**E)** A plot of the potentiation of the 2^nd^ alkaline transient relative to the 1^st^, across different axon firing frequencies and AZ release frequencies. The vertical dotted line represents the *average* firing frequency for this motor neuron.

## DISCUSSION

This study was driven by the cognitive dissonance created by recent imaging data showing alkalinization without a trace of acidification at conventional glutamatergic synapses [14], despite numerous studies that have demonstrated ASIC activation at conventional glutamatergic synapses. Our computational approach addresses this paradox and makes the following conclusions:

i. A degree of acidification is inevitable at the site of exocytosis at conventional glutamate synapses, although the time course is ≪ 1 ms.
ii. Phosphate buffering has the capacity to almost entirely extinguish acidic transients at and above physiological levels (~1 mM in humans; unknown in *Drosophila*), but bicarbonate buffering has little impact on the acid spike.
iii. Assumptions regarding the degree to which SVs are filled with Glu, and the time course of fusion pore opening, have little impact on the acid spike. Interestingly, co-loading with ATP substantially enhances the acidic spike.
iv. PMCA-mediated cleft alkalinization is systematically diminished by increasing CA activity, as observed empirically, and although it is almost entirely extinguished by 1 mM phosphate in the open cleft, phosphate is far less influential in a closed cleft.
v. Cleft morphology, specifically, continuity with a neighboring void, is essential for controlling PMCA-mediated alkalinization, as cleft pH homeostasis will “fail” below a certain level of CA activity. The ability of CO_2_ to cross membranes gives greater power to bicarbonate buffering over phosphate buffering wherever cleft volume is restricted.
vi. Conditions for the summation of alkaline transients at individual AZs, from one AP to the next, would require either higher axon firing frequencies than those observed to date, or higher AZ release probabilities.

Ultimately the paradox is resolved, as it was previously anticipated [47] that “clefts harbor exceedingly rapid acidic transients at the level of microdomains”, but that “detecting such rapid events with pH imaging would require sampling rates of ~10 kHz and knowledge of when release occurs at a particular release site to allow signal averaging.”

The claim that acidification is inevitable at the site of exocytosis requires some qualification. The *Drosophila* NMJ cleft is exceedingly narrow (13.5 nm), but even if the distance was doubled [it is 28 ± 9 nm at the rat Calyx of Held [48]] the consequence is minimal, as our model still shows cleft equilibration with SV pH (5.5) but with ½-width of acidification somewhat diminished (from 0.123 to 0.080 ms). Phosphate buffering has the greatest impact on the acid spike, with 1 mM total phosphate halving it’s depth and greatly reducing its ½-width (from 0.123 to 0.006 ms). It should be noted that HPO_4_^2−^ is only present at 0.5 mM in our model, with H_2_PO_4_^−^ present at 0.5 mM. This can be put into context with inorganic phosphate levels in our blood (~ 1 mM), which confers phosphate buffering of ~0.8 mM HPO_4_^2−^ and ~0.2 mM H_2_PO_4_^−^, when adjusted for a pH of 7.4 rather than 7.2 used here. Considering the activity coefficients of these ions, as much as 4 mM phosphate would need to be added to a physiological saline such as ours to accomplish this sort of phosphate buffering power [49]. We should emphasize that the open boundary condition used here, i.e. access to an infinite buffer reservoir at a distance of 0.4 μm, is ideal, whereas it is likely that there will be greater impediment to exchange at clefts *in vivo*. Certainly, the other ideal boundary condition of a closed cleft greatly curtails the ability of phosphate to dampen alkalinization. To the extent that an acid spike is not extinguished by the buffering system, an acid spike on the order of only 0.1-0.5 ms is well able to activate ASICs [43, 44]. The capacity of phosphate buffering to tune transduction between SV exocytosis and ASIC activation therefore provides an intriguing window into a possible mechanism of action linking either hypophosphatemia or hyperphosphatemia with associated neurological disorders [50, 51].

While the likely presence of acidic microdomains can resolve the paradox, it is the alkaline signal that captured our attention, as it’s integral is ~ 3 orders of magnitude larger. Unlike the acid transient, the slow alkaline transient is likely to persist from one AP to the next during a train of APs. However, as pointed out in **Figure 6**, this impression is misleading, as empirical data collected from groups of AZs (≫9) do not represent pH change at individual AZs that release with an average probability of 0.11. The axon would have to fire at a frequency of 40 Hz or greater (double the axon’s native firing frequency of 21 Hz) before the “average” AZ would release at a rate where alkaline transients persisted from one release event to the next. The significance of transients persisting is that it allows for the summation of pH change at an AZ, thus providing a mechanism for activity-dependent potentiation of release through the pH sensitivity of exposed synaptic proteins. Nevertheless, for the following reasons we are not ready to relinquish the possibility of such a gain mechanism. First, as at many other NMJs, release probability at the *Drosophila* NMJ is quite heterogeneous between AZs [52, 53], in which case any AZ with a probability twice the average, e.g. 0.22, could be potentiated at frequencies marginally over the native firing frequency. Secondly, the native firing frequency was quantified in restrained animals where the fictive locomotion representing peristaltic contractions was drawn out ten-fold, raising the question of whether firing frequency during each contraction was underestimated relative to a freely-behaving animal. Lastly, the model was only extrapolated to examine the potentiation of a second release event relative to the first – due to concerns about extrapolating the model well beyond assumptions made for individual release events. It seems reasonable to think that alkaline transient summation might proceed in a supralinear fashion as buffers deplete faster than they can be replenished in the synaptic cleft. Some support for the latter two points comes from the trans-cuticle imaging data of Stawarski et al., 2020, where cleft alkaline transients exceeded 1 log unit at some nerve terminals in the intact animal.

Perhaps the most surprising result was the finding of the necessity to “ventilate” the synapse to allow neurotransmitters to escape and exchange buffering entities with a larger volume. Part of the effect was likely a consequence of limitations of the model, where the movement of species beyond Ca^2+^, H^+^ and CO_2_ across membranes was not accommodated. For instance, the model did not allow for Glu reuptake or bicarbonate movement across membranes, and all proton movement across membranes was confined to exocytosis and the exchange of 2H^+^ for Ca^2+^ by PMCA. Despite these shortcomings, it seems unlikely that the depth of the pH spike would have been ameliorated by further sophistication of the model, such was the speed of the phenomenon. Furthermore, although the alkaline transient was primarily the product of PMCA activity, and heavily dependent on the level of CA activity, the effect of closing access to the voids of the SSR was robust. Both acidification and alkalinization events highlight the dynamic pH microenvironment within nervous tissue, and the limitations of pH homeostasis. Just as there are limitations to Ca^2+^ entry from the synaptic cleft [54], the density of synapses and their morphology might be limited by a need to accommodate pH transients and pH homeostatic mechanisms.

## MATERIALS AND METHODS

### Experimental

#### pH imaging and Immunohistochemistry

The experimental techniques and equipment used here were identical to those used by [14]. The fluorescent pH reporter pHusion-Ex was expressed in muscle fibers using the 24B-GAL4 [55] (**Fig.1A**). pHusion-Ex consists of superecliptic pHluorin with an N-terminal IgG κ secretion signal linked to FusionRed with a C-terminal transmembrane sequence from the human platelet-derived growth factor receptor (hPDGFR) [14]. Alexa Fluor 647 dextran 10,000 MW (AF647; D22914, Life Technologies) was forward-filled into motor nerve terminals using the technique described in [56] (**Fig. 1 B**). Female *Drosophila* 3^rd^ instar larvae were fillet dissected in Schneider’s insect medium and transferred to HL6 physiological solution for live imaging [57]. Nerves to the hemisegment of interest were not cut except when an axon was forward-filled. Images of the live NMJ were collected on a Nikon AR1 laser-scanning confocal microscope using a 60 × 1.2 NA water-immersion apochromat Nikon objective (**Fig. 1B**). This confocal, and the wide-field imaging system described below, were controlled by Nikon NIS-Elements software. 16-bit images were subsequently converted to 8-bit for assembly into figures using Canvas X software (ACD Systems).

Changes in pHusion-Ex fluorescence (ratio) were calculated from pairs of images collected on a Nikon Eclipse FN1 microscope fitted with a 100 × 1.1 NA water-immersion objective using an Andor TuCam beam-splitter to direct emission wavelengths to two separate Andor iXon3 EMCCD cameras. Fluorophores were excited using a SPECTRA-X light source and images were collected at rates of either 94 pairs of images per second (**Fig. 1C**), or 10 pairs of images per second (**Fig. 1D**). Images were analyzed offline using ImageJ and the fluorescence ratio of pHusion-Ex was compared to the calibration of Stawarski et al., (2020) to yield values of pH change.

For the purposes of immunohistochemistry, fillet dissected preparations were fixed with Bouins solution for 1 minute before subsequent processing as described by Stawarski et al. 2020. The following antibodies were used: 1°: nc82 (1:200) (mouse) from DSHB Hybridoma Bank (supernatant) and rabbit anti-PMCA, a gift from Dr. Gregory Lnenicka at SUNY Albany (1:200); 2°: goat anti-mouse AF488 (1:500) (Invitrogen, A-11001), and donkey anti-rabbit Cy5 (1:200) (Jackson ImmunoResearch). Preparations were mounted under glass coverslips in SlowFade Gold (Invitrogen, S36937) and imaged using a 100 × 1.3 NA Olympus oil objective.

### Electron microscopy

Female *Drosophila* 3^rd^ instar larvae were filet dissected and fixed for 1 hr with ice cold 3% paraformaldehyde and 2.5% glutaraldehyde made up in 100 mM Sørensen’s phosphate buffer (PB; pH 7.4) containing 2 mM CaCl_2_. Following fixation, larvae were washed with PB 5 × 2 mins. Any remaining fixative was then quenched with a 20 mM glycine solution for 5 mins on ice, then washed with PB 3 × 2 minutes. Larvae were incubated in 1% aqueous OsO_4_ for 30 mins at room temperature (RT), then the solution was replaced with aqueous 1.5% potassium ferrocyanide for another 30 mins in the dark at RT. Larvae were washed with water 3 × 10 mins. Larvae were incubated in aqueous 1% uranyl acetate for 1 hr in the dark at RT, then washed. Larvae were dehydrated in an ascending ethanol series for 10 mins each (30%, 50%, 70%, 90%, 100%, 100%), then 1:1 ethanol to acetone for 5 mins and 100% acetone for 2 × 10 mins. For resin infiltration, larvae were placed in a 3:1 solution of acetone to Durcupan ACM resin mixture (Sigma-Aldrich) for 2 hrs, 1:1 acetone to resin for 2 hrs, and 1:3 acetone to resin overnight. Larvae were placed in 100% resin for 4 hrs under vacuum, then flat embedded between slide glass and an aclar sheet (Electron Microscopy Sciences) and polymerized at 60 °C for 2 days. Embedded larvae were mounted on resin blocks, then sectioned at 50 nm and collected on TEM grids. Samples were observed with a Tecnai G2 Spirit transmission electron microscope (Thermo Fisher Scientific) at 100 kV, and images were acquired using a Veleta CCD camera (Olympus) operated by TIA software (Thermo Fisher Scientific). Cleft widths were measured from 12 active zones (AZs) across 5 NMJs from muscle 6/7, 13 and 12. Measurements were taken every 30 nm along each AZ and measured as the distance between the outer leaflet of the bouton plasma membrane and the outer leaflet of the SSR plasma membrane.

### Theoretical

#### Sections i-iv below address different aspects of the computational model

##### i) Geometry

Assuming, for simplicity that AZs form a square lattice with regular periodicity in the *x* and *y* axis, we can restrict the domain of the numerical calculations to a “primitive” cell of the square lattice and impose periodic boundary conditions on the concentration of all the species, c_i_(x, y, z, t) **Table 1**. The periodic boundary conditions dictate that c_i_(x, y, z, t) = c_i_(x + m × L, y + n × L, z, t), where m and n are integers and L is the distance between the centers of two primitive cells along either the x or y axis. The height of the primitive cell (13.5 nm) was determined to be substantially less than its lateral extent (800 nm) (**Fig. 2C-H**), allowing us to neglect the *z* dependence of the concentration fields, and treat *Eq. 1* (see Results) as a two-dimensional problem. In our numerical computations, we impose two distinct boundary conditions. The first, based on our observations from electron microscopy (EM) images, where each primitive cell is connected to wide channels at its outer edges (**Fig. 2B-D**), informs a Dirichlet boundary condition. These channels may be viewed as an infinite reservoir of buffers, which implies that the concentration of all the species at the edges of the AZs may be taken to be constant throughout neuronal activity, i.e., c_i_(r = R, t) = const. The second boundary condition, represents a Neumann boundary condition, where we assume that there is no net flux of any species at the edge of the primitive cell; which implies that at the edges of AZs, [dc_i_(r, t)/dr]_r=L_ = 0. Although there is no way to accurately determine the correct boundary conditions for the cleft, we believe that the Dirichlet boundary condition is more representative of the *Drosophila* NMJ. Indeed, as we show below, the results of our simulation using the Dirichlet boundary condition are largely in agreement with the experimental data. However, applying the Neumann boundary condition shows that the pH does not settle to the baseline in the time scale observed in the experiments unless we assume the CO_2_ concentration is constant in the cleft (ideal scenario where CO_2_ diffuses freely across membranes bounding the cleft) and the facilitation by carbonic anhydrase (CA) is above a certain threshold. Lastly, for both boundary conditions we assume azimuthal symmetry about the release site, which we take to be the origin of the primitive cell. Therefore, the concentration fields are a function of the radial distance away from the origin. In effect, the problem was reduced from a three-dimensional to a one-dimensional problem. To preserve the area of the primitive cell, we take its radius to be 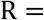 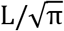, which gives the same area/volume for the primitive cell. Note that we assume azimuthal symmetry to simplify numerical calculations but verified that errors coming from edge effects are negligible.

##### ii) Diffusion

We start with the diffusion term of *Eq. 1* for species *i* and write it in polar coordinates, with the assumption of azimuthal symmetry as:

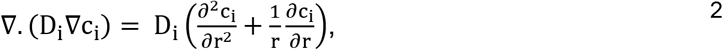

where the diffusion coefficient is assumed to be constant throughout the cleft. Each of species 1-12 is identified in **Table 1**.

To implement our model numerically, we first employ the method of lines to discretize Eq. 2 in space and then divide the area of interest ( 0 ≤ r ≤ R) into n concentric circles. Here, the value of n is chosen with the purpose of being able to run the simulation within a reasonable time (10-15 minutes, n = 57). To apply the boundary conditions numerically, we use the same method as that described in [31, 58] for both innermost (the release point) and outermost shell (the edge of primitive cell), with the exception that the outermost shell has two possibilities as discussed in the geometry section. The details of the finite element discretization and boundary conditions for the numerical calculations can be found in the Appendix. All the species are assumed to be initially at equilibrium (just before SV exocytosis) and are homogeneously distributed in the cleft. The initial values for the concentrations can be found in **Table 3**, for details on calculations see the buffers section. Diffusion coefficients, listed in **Table 2** are measured in water, but in our simulations we use the effective diffusion coefficient ( D* = D/γ^2^) where γ is the tortuosity of the environment [59], and is assumed to be 1.5 for the synaptic cleft [60–62].

**Table 2.** (Kinetic rates, diffusion coefficients, and assorted constants)

| Parameter | Value | Source | Parameter | Value | Source |
| --- | --- | --- | --- | --- | --- |
| $k_{1,2}$ | $4.15 * 10^{16} \frac{1}{mM.s}$ | [31] | $k_{13,12}$ | $10^{10} \frac{1}{s}$ | Chosen to achieve equilibrium |
| $k_{2,1}$ | $1 * 10^{16} \frac{1}{s}$ | Chosen to achieve equilibrium | $D_{H^+}$ | $8.69 * 10^{-5} \frac{cm^2}{s}$ | [31] |
| $k_{2,3}$ | $10.9631 \frac{1}{s}$ | [31] | $D_{HCO_3^-}$ | $1.11 * 10^{-5} \frac{cm^2}{s}$ | [31] |
| $k_{3,2}$ | $0.0302 \frac{1}{s}$ | [31] | $D_{H_2CO_3}$ | $1.11 * 10^{-5} \frac{cm^2}{s}$ | [31] |
| $k_{4,5}$ | $1.584 * 10^{14} \frac{1}{mM.s}$ | [79] | $D_{CO_2}$ | $1.71 * 10^{-5} \frac{cm^2}{s}$ | [31] |
| $k_{5,4}$ | $1 * 10^{10} \frac{1}{s}$ | Chosen to achieve equilibrium | $D_{H_2PO_4^-}$ | $1.56 * 10^{-5} \frac{cm^2}{s}$ | [31] |
| $\frac{k_{7,6}}{k_{6,7}}$ | $5.62 * 10^{-2} \frac{1}{mM}$ | [80] | $D_{HPO_4^{2-}}$ | $1.56 * 10^{-5} \frac{cm^2}{s}$ | [31] |
| $\frac{k_{9,8}}{k_{8,9}}$ | $2.14 * 10^{-7} mM$ | [80] | $D_{Glu}$ | $0.76 * 10^{-5} \frac{cm^2}{s}$ | [81] |
| $En_{acc}$ | 600 | Adjusted | $D_{Glu^-}$ | $0.76 * 10^{-5} \frac{cm^2}{s}$ | [81] |
| $k_{6,7}$ | $10^{10} \frac{1}{s}$ | Chosen to achieve equilibrium | $D_{\text{Glu}^{2-}}$ | $0.76 * 10^{-5} \frac{\text{cm}^2}{s}$ | [81] |
| $k_{8,9}$ | $10^{10} \frac{1}{s}$ | Chosen to achieve equilibrium | $D_{\text{ATP}^{4-}}$ | $0.71 * 10^{-5} \frac{\text{cm}^2}{s}$ | [82] |
| $\frac{k_{11,10}}{k_{10,11}}$ | $3.412 * 10^{-4} \text{ mM}$ | [77] | $D_{\text{ATP}^{3-}}$ | $0.71 * 10^{-5} \frac{\text{cm}^2}{s}$ | [82] |
| $k_{11,10}$ | $10^{10} \frac{1}{s}$ | Chosen to achieve equilibrium | $D_{\text{ATP}^{2-}}$ | $0.71 * 10^{-5} \frac{\text{cm}^2}{s}$ | [82] |
| $\frac{k_{12,13}}{k_{13,12}}$ | $\frac{1}{\text{mM}}$ | [77] | $\lambda$ | 1.5 | [62] |

##### iii) Buffers

The second term in *Eq. 1* represents chemical reactions of all the species in our model, i.e., buffering, which plays a vital part in the pH regulation [1, 63, 64]. We consider the following reactions for bicarbonate, phosphate, glutamate (Glu), and ATP:

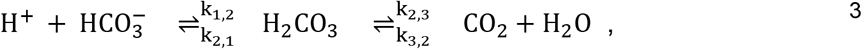

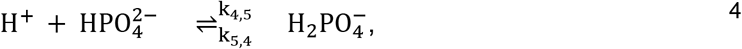

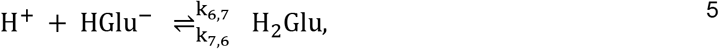

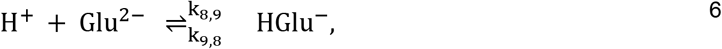

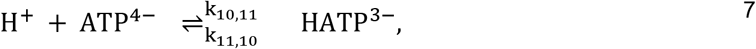

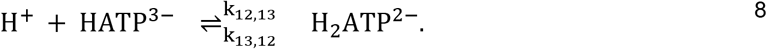

Where *Eq. 3* and *4* denote the bicarbonate and phosphate buffering reactions, respectively. *Eq. 5* and *6* describe the protonation of different species of Glu and *Eq. 7* and *8* describe protonation of different species of ATP. We note that all reactions, except for the hydration/dehydration of CO_2_ in *Eq. 3*, are extremely fast when compared to the time scale of the diffusion of all the species [31]. Here, with the exception of the hydration/dehydration of CO_2_ in *Eq. 3,* we choose the rates of reactions from *Eqs. 3–8* to be as fast as possible with the result that the concentration dynamics of all the species are limited by diffusion [22, 31, 65, 66]. The rates of these reactions can be found in **Table 2**. Note that we have ignored the reaction 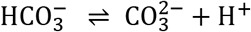 (which is related to Eq. 3), because it occurs at a high pH well outside the pH range of the experiments we are modeling. Furthermore, the reaction H_2_CO_3_ ⇌ CO_2_ + H_2_O is normally very slow in comparison to the rest of the reactions, but by adding CA enzymatic activity, the reaction gets faster and may render this reaction relevant. Rather than including it in our reaction diffusion equations, we simply implement this facilitation by augmenting the rate of hydration/dehydration of CO_2_ to a larger value (En_acc_) [31, 59, 67], as found in **Table 2**. Thus, we write the mass reaction rate equations as:

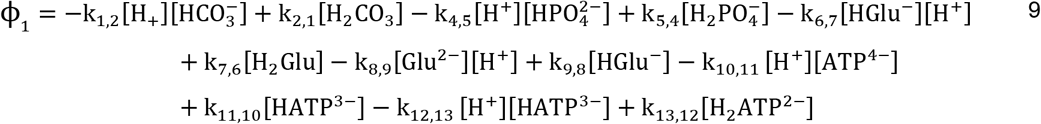

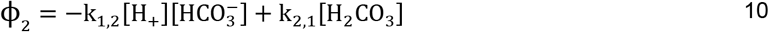

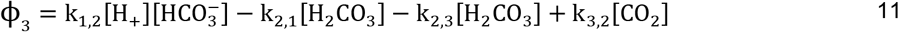

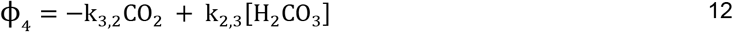

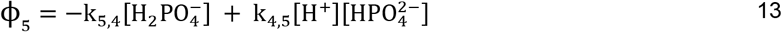

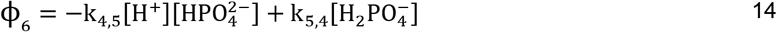

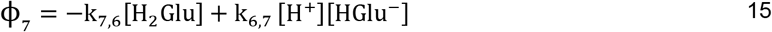

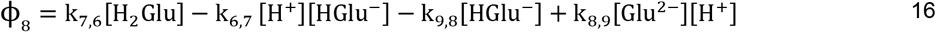

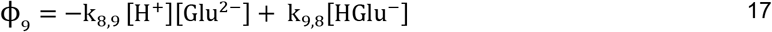

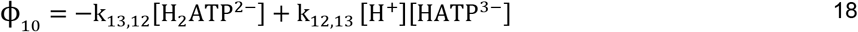

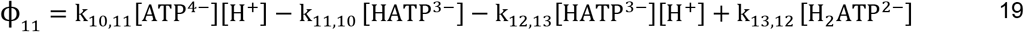

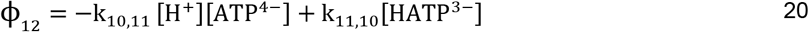

To calculate the initial concentrations of the species above, which can be found in **Table 3**, we simply assume that all the species are at an equilibrium before the AP (ϕ_i_ = 0). Applying this equilibrium condition to *Eq. 9*–*20* allows us to obtain the initial concentration of each of the species. Because the extracellular concentrations of Glu and ATP are in the *μM* range [68, 69], they do not play a significant role in buffering and we assume their steady state concentrations to be zero.

**Table 3.**
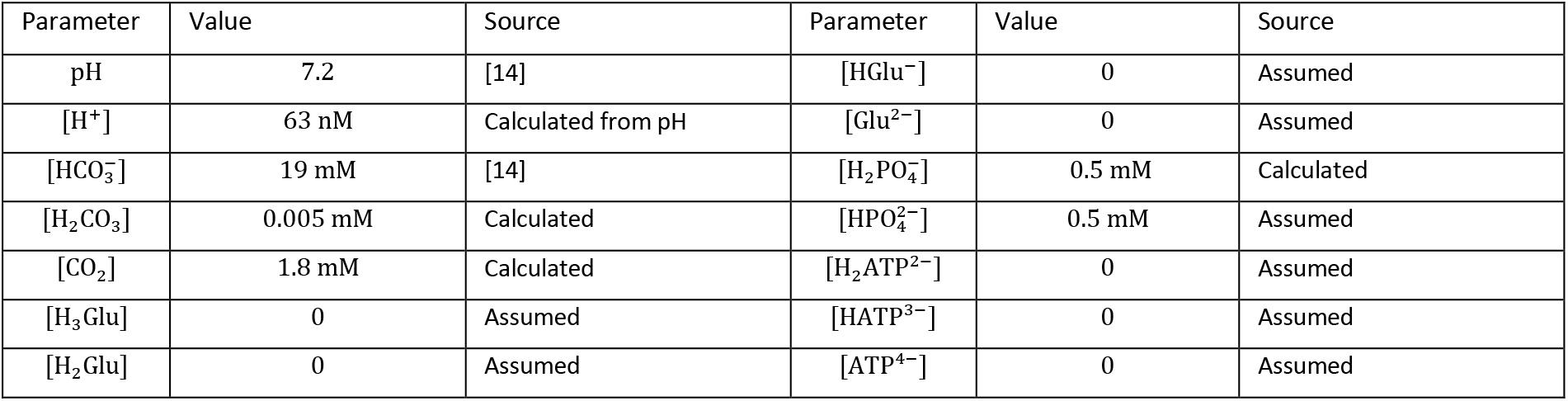
(Initial and steady-state concentrations in the cleft)

| Parameter | Value | Source | Parameter | Value | Source |
| --- | --- | --- | --- | --- | --- |
| pH | 7.2 | [14] | $[\text{HGl}u^-]$ | 0 | Assumed |
| $[\text{H}^+]$ | 63 nM | Calculated from pH | $[\text{Glu}^{2-}]$ | 0 | Assumed |
| $[\text{HCO}_3^-]$ | 19 mM | [14] | $[\text{H}_2\text{PO}_4^-]$ | 0.5 mM | Calculated |
| $[\text{H}_2\text{CO}_3]$ | 0.005 mM | Calculated | $[\text{HPO}_4^{2-}]$ | 0.5 mM | Assumed |
| $[\text{CO}_2]$ | 1.8 mM | Calculated | $[\text{H}_2\text{ATP}^{2-}]$ | 0 | Assumed |
| $[\text{H}_3\text{Glu}]$ | 0 | Assumed | $[\text{HATP}^{3-}]$ | 0 | Assumed |
| $[\text{H}_2\text{Glu}]$ | 0 | Assumed | $[\text{ATP}^{4-}]$ | 0 | Assumed |

##### iv) Extrusion

The third term in *Eq. 1*, *J_i_*, denotes the extrusion of H^+^ from the cleft through the PMCA which exchanges Ca^2+^ for 2H^+^. Note that the only non-zero term, *J*_1_, is calculated from a kinetic model of the PMCA based on the Michaelis-Menten equation. Although several H^+^ exchangers such as Na^+^/H^+^ and Cl^−^/H^+^ on the plasma membrane have the potential to remove H^+^ from the cleft, it is the PMCA activity which is responsible for cleft alkalinization [14, 20]. As previously mentioned, PMCA is vital for the clearance of Ca^2+^ from both the pre-and postsynaptic compartments during synaptic activity [32, 33]. Most of the Ca^2+^ which enters the cytosolic compartment is bound to mobile and immobile Ca^2+^ buffers, but subsequently intercepted by the PMCA, Na^+^/Ca^2+^ exchanger (NCX), mitochondria and the endoplasmic reticulum (ER); the latter two acting as Ca^2+^ traps. In an electroneutral process, the PMCA exchanges one Ca^2+^ for 2H^+^ in the cleft, thereby driving pH changes in the cleft [70]. To model this process at pre- and postsynaptic compartments, we make use of the Michaelis–Menten kinetics for a one-substrate enzyme-catalyzed reaction, where PMCA is taken to be the enzyme and Ca^2+^ as the substrate. For simplicity, we assume that both the pre- and postsynaptic compartments are modeled with the same process, but with different densities of PMCA and peaks in Ca^2+^ concentration. Thus, we write the chemical reaction for this process as [71, 72]:

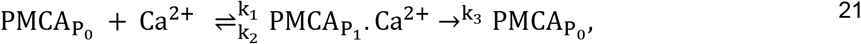

where PMCA can take on two conformational states denoted by P_0_ and P_1_, where P_0_ has a high affinity for Ca^2+^ and P_1_ has a low affinity for Ca^2+^. P_0_ is localized at the intracellular leaflet of both pre- and postsynaptic compartments while P_1_ is exposed to the cleft at the extracellular leaflet of both the pre- and postsynaptic compartments [71]. Note that there is a turnover between *P*_0_ and *P*_1_ with a rate of *k*_3_. Because PMCA is continually operating, even in the absence of an AP, there should be a steady leakage of Ca^2+^ into the pre- and postsynaptic compartments in order to achieve a steady state condition [73]. At steady state, we assume the rate at which the H^+^ are removed from the cleft by PMCA equals the rate of leakage of H^+^ from the pre- and postsynaptic compartment to the cleft. In other words, we neglect the baseline H^+^ extrusion from the cleft.

The Michaelis-Menten equation gives the rate at which H^+^ are being extruded from the cleft by PMCAs [73]:

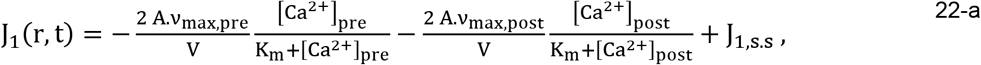

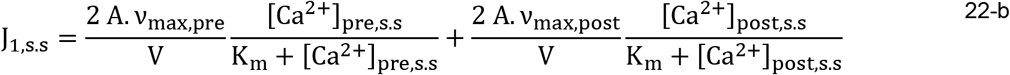

where K_m_ = (k_2_ + k_3_)/k_1_ is the half-maximal activating concentration, ν_max,pre_ = n_pre_k_3_ and ν_max,post_ = n_post_k_3_ are the maximal velocity of transport 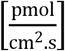 in pre- and postsynaptic compartments, respectively, n_pre_ and n_post_ are the density of PMCA at pre- and postsynaptic compartment, respectively, A is the area, and V the volume of the primitive cell. In Eq. 22-b [Ca^2+^]_pre,s.s_ and [Ca^2+^]_post,s.s_ are steady-state concentrations of Ca^2+^ at pre- and post-synaptic compartments, respectively and J_1,s.s_, which is a constant, is the steady-state extrusion of H^+^ from the cleft. Note also that J_1_ has no spatial dependence as we assume that the PMCA is distributed uniformly on pre- and postsynaptic compartments and that Ca^2+^ is homogeneously distributed in the cleft. Note that in Eq. 22-a, [Ca^2+^]_pre_ and [Ca^2+^]_post_ are concentration of Ca^2+^ at pre- and postsynaptic compartments, respectively at each time and the “2” accounts for the fact that 2H^+^ exchange for Ca^2+^. Furthermore, 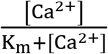 can be interpreted as the fraction of PMCA on pre- and postsynaptic compartments that would exchange 2H^+^ with Ca^2+^ to the cleft at time *t*.

To proceed, both [Ca^2+^]_pre_ and [Ca^2+^]_post_ are required as a function of time. Ideally, we would implement Ca^2+^ as a separate species in our reaction-diffusion equation along with all its buffers and organelles that take up Ca^2+^ and release it in an intermittent fashion on pre- and postsynaptic compartment. However, this would make the model unreasonably complicated for the following three reasons: 1) complex geometry of the postsynaptic compartment, 2) lack of information on the location and density of Ca^2+^ permeable channels, 3) the complexity of mechanisms which regulate intracellular Ca^2+^ concentration that needs the implementation of Monte Carlo methods. For simplicity, we postulate that both [Ca^2+^]_pre_ and [Ca^2+^]_post_ take the following form as a function of time:

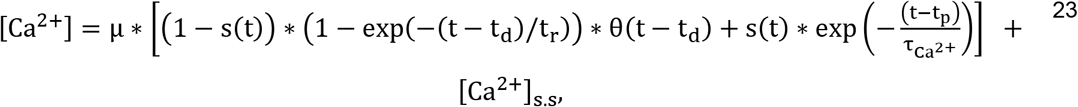

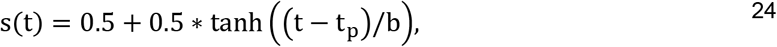

Where, θ(t) is the step function, and t_d_, t_r_, t_p_, 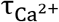 and μ are: latent time (t < td, [Ca^2+^] = steady state Ca^2+^ concentration), tau of rise, time of peak, tau of decay, and maximum Ca^2+^ concentration, respectively. It is important to note that these parameters take on different values on the pre- and postsynaptic compartments of the cleft. This is because different Ca^2+^ entry and Ca^2+^ regulation mechanisms operate on either side. Therefore, [Ca^2+^]_pre_ and [Ca^2+^]_post_ should have different kinetics and time courses. For example, in the presynaptic compartment, Ca^2+^ entry is through VGCCs, while on postsynaptic compartment it is through GluRs. This causes t_p_ of the presynaptic compartment to be much shorter than that of the postsynaptic compartment. The values of t_d_, t_p_, t_r_, 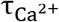 and *μ* for the presynaptic compartment are approximated from [45], while for the postsynaptic compartment t_d_, t_p_, t_r_ are approximated from measurements made from two-electrode voltage clamp (TEVC) recordings in our laboratory (data not shown). The value for 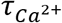 in the postsynaptic compartment is obtained from [32] and *μ* is adjusted to capture experimental results (for more details, see the numerical simulation section). All the parameters of PMCA and Ca^2+^ concentration can be found in **Table 5**. As depicted in **Figure S1**, the time course of Eq. 23 is such that for t < t_d_, [Ca^2+^] is at the steady state Ca^2+^ concentration and then rises quickly to reach a peak between time t_d_ < t < t_p_, then for t > t_p_ the concentration decays exponentially. The features of these time courses are generally reflected in data from Ca^2+^ imaging experiments [32, 45, 57].

**Table 4.** (Source parameters in the SV lumen)

| Parameter | Value | Source | Parameter | Value | Source |
| --- | --- | --- | --- | --- | --- |
| pH | 5.5 | [76] | $N_{0,\text{glutamate}}$ | 8000 | [75] |
| $[\text{H}^+]_{\text{free}}$ | 3.2 $\mu\text{M}$ | Calculated | $[\text{H}_2\text{Glu}]_{\text{SV}}$ | 69.97mM | Calculated |
| $\tau_f$ | 1 $\mu\text{s}$ | Assumed [41] | $[\text{HGl}u^-]_{\text{SV}}$ | 1242.9 mM | Calculated |
| $r_{\text{SV}}$ | 13.42 nm | [83] | $[\text{Glu}^{2-}]_{\text{SV}}$ | 0 | Calculated |
| $N_{0,\text{ATP}}$ | 610 | Assumed (to arrive at 100 mM total ATP in SV) | $[\text{HATP}^-]_{\text{SV}}$ | 87.28 mM | Calculated |
| $[\text{H}_2\text{ATP}]_{\text{SV}}$ | 3.47 mM | Calculated | $[\text{ATP}^{2-}]_{\text{SV}}$ | 9.35 mM | Calculated |

**Table 5.** (Extrusion parameters)

| Parameter | Value | Source | Parameter | Value | Source |
| --- | --- | --- | --- | --- | --- |
| $K_m$ | 0.83 $\mu\text{M}$ | Adjusted [72, 73, 84] | $k_3$ | 100 $\frac{1}{\text{s}}$ | [72] |
| $n_{\text{pre}}$ | 150 $\frac{1}{\mu\text{m}^2}$ | Adjusted (from glutamate experiment [14]) | $n_{\text{post}}$ | 1750 $\frac{1}{\mu\text{m}^2}$ | Adjusted |
| $\mu_{\text{pre}}$ | 0.4 $\mu\text{M}$ | [45] | $\mu_{\text{post}}$ | 2.5 $\mu\text{M}$ | Adjusted |
| $t_{d,\text{pre}}$ | 0.5 ms | [85] | $t_{d,\text{post}}$ | 0.9 ms | [85] |
| $t_{p,\text{pre}}$ | 1.5 ms | [85] | $t_{p,\text{post}}$ | 39.3 ms | Our TEVC experiments |
| $t_{r,\text{pre}}$ | 0.2 ms | Adjusted | $t_{r,\text{post}}$ | 8 ms | Adjusted |
| $b_{\text{pre}}$ | 0.5 ms | Adjusted [45] | $b_{\text{post}}$ | 10 ms | Adjusted |
| $\tau_{\text{Ca}^{2+},\text{pre}}$ | 50 ms | [45] | $\tau_{\text{Ca}^{2+},\text{post}}$ | 50 ms | [32] |
| $\mu_{\text{pre(train)}}$ | 1.2 $\mu\text{M}$ | [32] | $t_{r,\text{pre (train)}}$ | 27 ms | Adjusted [57] |
| $t_{d,\text{pre(train)}}$ | 0.5 ms | Adjusted [45] | $b_{\text{pre(train)}}$ | 0.5 ms | Adjusted [45, 57] |
| $t_{p,\text{pre(train)}}$ | $3 * \left(\frac{1000}{21}\right) \text{ms}$ | [57] | $\tau_{\text{Ca}^{2+},\text{pre(train)}}$ | 68 ms | [57] |

##### v) Source Terms

The last term in Eq. 1, S_i_, represents the source term of each species. The only non-zero terms of S_i_ are i = 1,7,8,9,10,11,12. In our model, we assume each AZ releases only one SV in addition to a complete fusion of the SV to the plasma membrane, i.e., a SV releases all of its contents. We also assume SVs contain Glu, ATP, and free H^+^. For the kinetics of release of Glu and ATP from the SVs, the number of molecules of each one of these three species in SVs at any time has the form 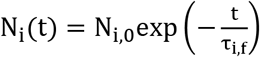, where τ_i,f_ is the time constant of fusion and release, and N_i,0_ is the total number of molecules of species i in a SV before the release. Therefore, 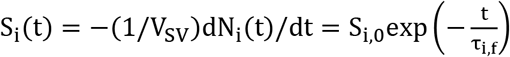, where V_SV_ is the volume of a SV and 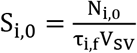. Previous measurements from high-resolution TEVC recordings of miniature excitatory junctional currents (mEJCs) suggest that the rise time of a mEJC is less than 100μs [41]. Therefore, most of the Monte Carlo models that use the kinetics of SV release assume the release of Glu and ATP to be instantaneous [72]. Here, we will also use a very small value as τ_i,f_ = 1 μs, except where indicated otherwise.

Given that N_i,0_ is dependent on the pH of the SV lumen, for the species i = 1, we calculate it by using the pH of the SV (pH 5.5). Whereas, for i = 7,8,9 and i = 10,11,12, which represent the different Glu and ATP species, respectively, we use titration plots (**Fig. S2**). In aqueous solutions, glutamic acid has four different charged and uncharged variants denoted by H_3_Glu^+^, H_2_Glu, HGlu^−^, Glu^2−^. In our numerical model, we ignore H_3_Glu^+^ because this variant only becomes relevant when pH is lower than 4.5-5 [74]. If we assume that before release, the glutamic acid molecules in a SV are at equilibrium, their concentrations can be obtained using *Eqs. 15, 16 and 17*, by setting ϕ_i_ = 0, with the constraint that *T*, the total number of Glu molecules (including all its variants) divided by the volume of SV, must be a constant.

Therefore,

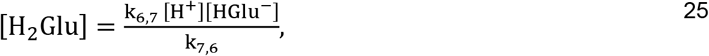

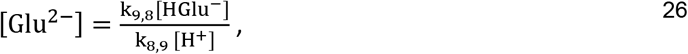

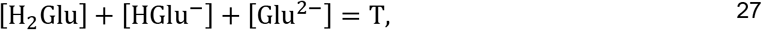

and therefore, we find

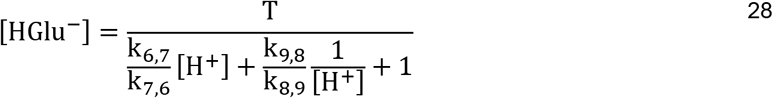

For example, for i = 7, 8, 9 if we assume to have a total of 8000 Glu molecules [75] at a SV pH of 5.5 [76], we have N_7,0_ ≈ 426, N_8,0_ ≈ 7574, N_9,0_ =0 (426 H_2_Glu, 7574 HGlu^−^, 0 Glu^2−^).

For i = 10,11,12, we can apply the same calculations as done for Glu above, to obtain the proportion of each species of ATP at different pH (**Fig. S2B**). The acid-base behavior of ATP has been studied in [77] and assume the ionic strength to be 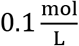.

Next, we require the total number of ATP molecules in SVs. Unfortunately, there is no information in the literature about the number of ATP molecules in a single glutaminergic SV. To solve this, we assume the number of ATP molecules is either zero, or that it is dictated by the osmotic pressure. This means that if a SV contains mostly ATP and Glu, the summation of total Glu concentration and ATP in the SV is equal to the osmotic pressure of the containing environment (~350 *mOsm*). As a result, one of our simulations is for the hypothetical scenario of 250 mM glutamate with 100 mM of ATP. Even though it seems that SVs can hold neurotransmitters above the cytosolic osmotic pressure [78], our assumption is only for simulating the effect of ATP on acid transients. It is worthwhile to mention that by comparing the titration figures of Glu and ATP, we observe that for the same concentration of both species, ATP will release substantially more H^+^ than Glu when both are released from an environment of pH 5.5 into an environment where they can equilibrate to a pH 7.2 (**Fig. S2A-B**).

##### vi) Numerical Simulation

To numerically solve *Eq. 1*, we first use the method of lines to discretize *Eq. 1* in space (semi-discretization). The origin of most values used are presented in **Table 5**. The resulting equations can be written in a matrix form with each element in the column vector representing the concentration of a particular species in each shell. After discretization, *n* stiff ordinary differential equations (ODE) remain, requiring special solvers [31]. To numerically solve the problem we used MATLAB’s ODE15s and a 16 core CPU – AMD Ryzen 3900X to run the code, where we set the relative and absolute tolerance to 10^−10^ to ensure scheme stability. Here, the output plots are restricted to pH vs time [31].

To test the utility of the model, we sought to test its numerical output against previously published empirical data [14]. However, to do so, we first had to recognize the mismatch between the model output, which generates cleft pH values for individual AZs, and the empirical data which represents a quantification of pHusion-Ex fluorescence collected from entire boutons harboring dozens of AZs, most of which do not release neurotransmitter with each AP. These fluorescence signals were compared to a calibration where all AZs around a bouton were simultaneously immersed in a saline series representing pH standards [14]. Such a protocol could provide the correct calibration for individual AZs only if all AZs did the same thing at the same time during an AP, i.e. all released neurotransmitter. A probability of release of anything less than 1 will lead to an underestimate of pH change at an individual AZ. Following in the footsteps of Stawarski et al. (2020), we examined the terminal of MN6/7-Ib which houses AZs with an average probability of release of 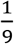 [45, 46]. If 9 AZs are found at each bouton then the empirical data represent the fluorescence response of 1 AZ firing at random, and this generates a ΔF/F signal of 1.94% [14]. As ΔF/F = (F_stim_−F_rest_)/F_rest_, we might recapitulate the fluorescence change at an individual AZ by dividing F_rest_ by 9. This is because the F_stim_ value represents an increase in fluorescence over an F_rest_ value which is approximately 9 fold smaller than the value of the entire bouton. The new ΔF/F signal of 17.5% can be compared directly with the calibration and yields ΔpH ≈ 0.225 for an individual AZ, equivalent to a change in pH from 7.200 to 7.425 for an individual AZ. Also see **Figure 6A-C**.

As mentioned throughout the manuscript, the values of some of the parameters such as the density of the PMCA, K_m_ of PMCA, Ca^2+^ concentration in postsynaptic microdomains where the PMCAs are located, and the concentration of ATP within glutamatergic SVs have eluded direct measurement at most synapses, not just at the *Drosophila* NMJ. For this reason, we drew on parameter values from multiple sources beyond the *Drosophila* NMJ, where all parameter values ultimately fell within a reasonable physiological range. Here we provide further explanation for a number of the parameters listed in **Table 5**. Starting with ν_max_, we note that a quantitative determination has been extremely difficult [72]. To get an estimated value of ν_max_ for *Eq. 22*, we require the values for k_3_, n_pre_ and n_post_. First, we assume that k_3_ on pre- and postsynaptic compartments are same with the value 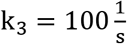 [72], which leads to ΔpH results consistent with our experimental data. If we use 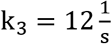 [84, 86], the resulting ΔpH would be significantly lower than the predicted experimental data. In addition, as discussed in [87], it is possible that the value of *k*_3_ measured *in vitro* is lower than the actual values of the PMCA *of in vivo* systems. Second, we need to know *n_pre_* and *n_post_*. Unfortunately, no direct measurements at the *Drosophila* NMJ have been reported to date, where reported values range from 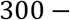 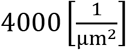 [87, 88]. To estimate *n_pre_*, we leveraged the experimental result where *7 mM* glutamic acid extinguishes alkalinization supplied by the postsynaptic compartment by desensitizing GluRs (**Fig. 2H** of [14]). Desensitization of GluRs prevents Ca^2+^ influx to the postsynaptic compartment. Therefore, as can be seen from Eq. 22, an absence of postsynaptic Ca^2+^ entry is synonymous with no removal of cleft H^+^ by postsynaptic PMCA activity. Any resulting alkaline transient is the product of presynaptic PMCA activity alone. However, presynaptic PMCA activity was insufficient to generate a cleft alkaline transient that could be distinguished from the noise (**Fig. 2H** of [14]), i.e. it would need to be less than ~0.0015 log units. It is also important to note that the presence of 7 mM glutamic acid is effective at buffering H^+^ near neutral pH, thus diminishing the amplitude of any alkaline transient generated by the presynaptic PMCA. Therefore, given that we are unable to detect a cleft alkaline transient solely from the presynaptic compartment as a consequence of the small signal in addition to the buffering of 7 mM glutamic acid, we used the noise of the data acquired from these experiments to obtain an approximate value of n_pre_. Here, we use ~0.0015 log units as maximum limit of alkalization rising from presynaptic for n_pre_. We observed that a value of 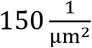 for n_pre_ replicated the empirical data, i.e. an alkaline transient no greater than 0.0015 pH units – in the presence of 7 mM glutamate (**Fig. S3**). A value of 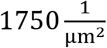 for n_post_ replicated the empirical data (**Fig. S3**). The corresponding values of ν_max_ for pre- and postsynaptic sides are 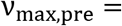 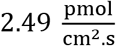 and 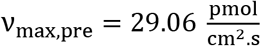 respectively, whereas the value of ν_max_ used in other models is in the range of 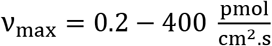 [73, 89–92]. Note: since PMCA activity obeys Michaelis-Menton kinetics and because the maximum ratio of activated PMCA is 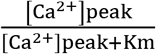, then for values of 0.4 μM Ca^2+^_pre_, 2.5 μM Ca^2+^_post_ and K_m_ = 0.83, the ratios for pre- and postsynaptic compartments will be 32% and 75%. Clearly, the difference in Ca^2+^ concentration in the two compartments cannot solely explain the empirical data (specifically, **Fig. 1C and Fig. S3**) and the method above was required to solve for unknown PMCA densities in the pre- and postsynaptic compartments. A third parameter requiring explanation, also relevant to Eq. 22, is the value of K_m_. As mentioned above, the expression, 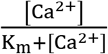, can be interpreted as the fraction of PMCA activated. However, if we choose a value of K_m_ in Eq. 22 lower than 0.5 μM, it leads to a flat peak in J_1_ for the postsynaptic compartment. Consequently, a flat peak for alkalinization in the results of the simulation. Therefore, the value of K_m_ is adjusted to a value of 0.8 μM, such that the kinetics and time course of ΔpH is consistent with our experimental data. The value of K_m_ used in other models ranges from 0.2 μM to 3 μM [73, 84, 87, 90].

To bring physiological context to the model, where neurons fire in bursts resulting in repeated release from the same AZs, we implemented a train of APs at 21Hz in our simulations. An individual AZ with release probability of 0.11 will fire at 21 × 0.11 = 2.3 Hz. The parameters of presynaptic Ca^2+^ concentration can be found in **Table 5**, where we have assumed it takes 4 nerve impulses to reach a plateau and remain constant until the last stimulus and then decay exponentially with τ = 68 ms. For simplicity, we have assumed that postsynaptic Ca^2+^ concentration for each AZ is the same as a single AP. Given that we are working with stiff ODEs, the results of the MATLAB’s ODE solvers do not have the same *dt*. In order to find the average of 9 AZs (**Fig. 6C**), we used MATLAB’s ‘pchip’ to interpolate between data points. After creating the same *dt* for all of the AZs, we found the average of all the AZs.

## Acknowledgements

This work was supported by NIH NINDS award NS103906 to GTM. Michal Stawarski’s current affiliation is with the *Department of Biomedicine, University of Basel, Basel, 4056, Switzerland*. We are grateful for discussions with Drs. Walter Boron, Robert Renden and Eugene Smith.

---

**Figure S1.**
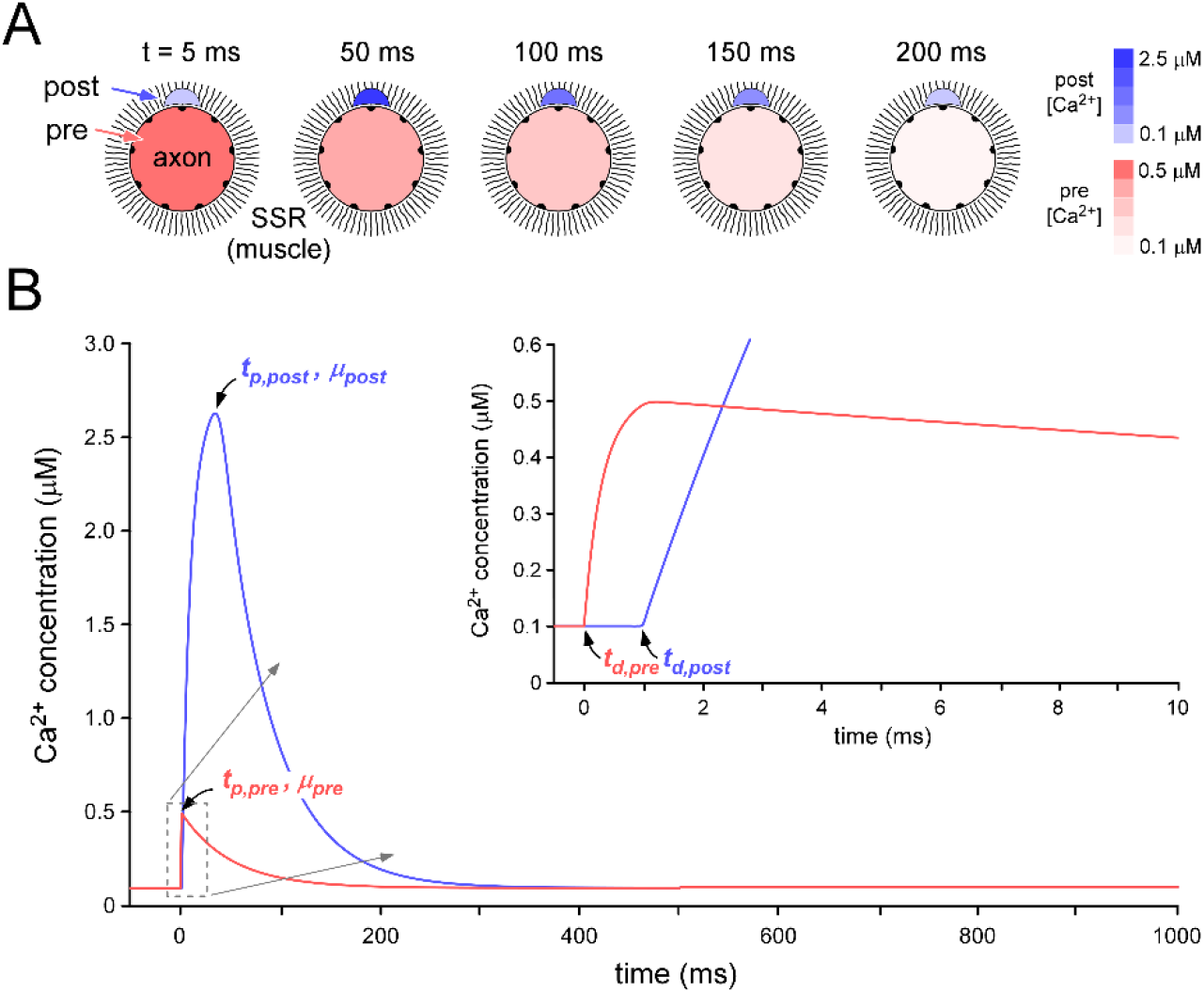
The time course of changes in Ca^2+^ concentration in pre- and post-synaptic compartments. (**A**) Stylized representations of changes in fluorescence of Ca^2+^ reporters on either side of the synaptic cleft responding to cytosolic Ca^2+^ changes associated with a pre-synaptic AP and the subsequent entry of Ca^2+^ into the postsynaptic muscle through GluR opposite a single presynaptic AZ. (**B**) Plots of the changes in cytosolic Ca^2+^ concentration over time on either side of the synapse ([Ca^2+^]_pre_ and [Ca^2+^]_post_). These plots differ due to the different mechanisms responsible for Ca^2+^ entry and Ca^2+^ extrusion on different sides of the synapse. Ca^2+^ enters the pre-synaptic compartment through VGCCs over a period of ~ 1ms during the later stages of the presynaptic AP. Ca^2+^ enters the postsynaptic compartment over a more extended time-course (~ 40 ms) through GluR after neurotransmitter is released from the pre-synaptic terminal. Inset; detail of the first 10 ms after the arrival of the pre-synaptic AP.

**Figure S2.**
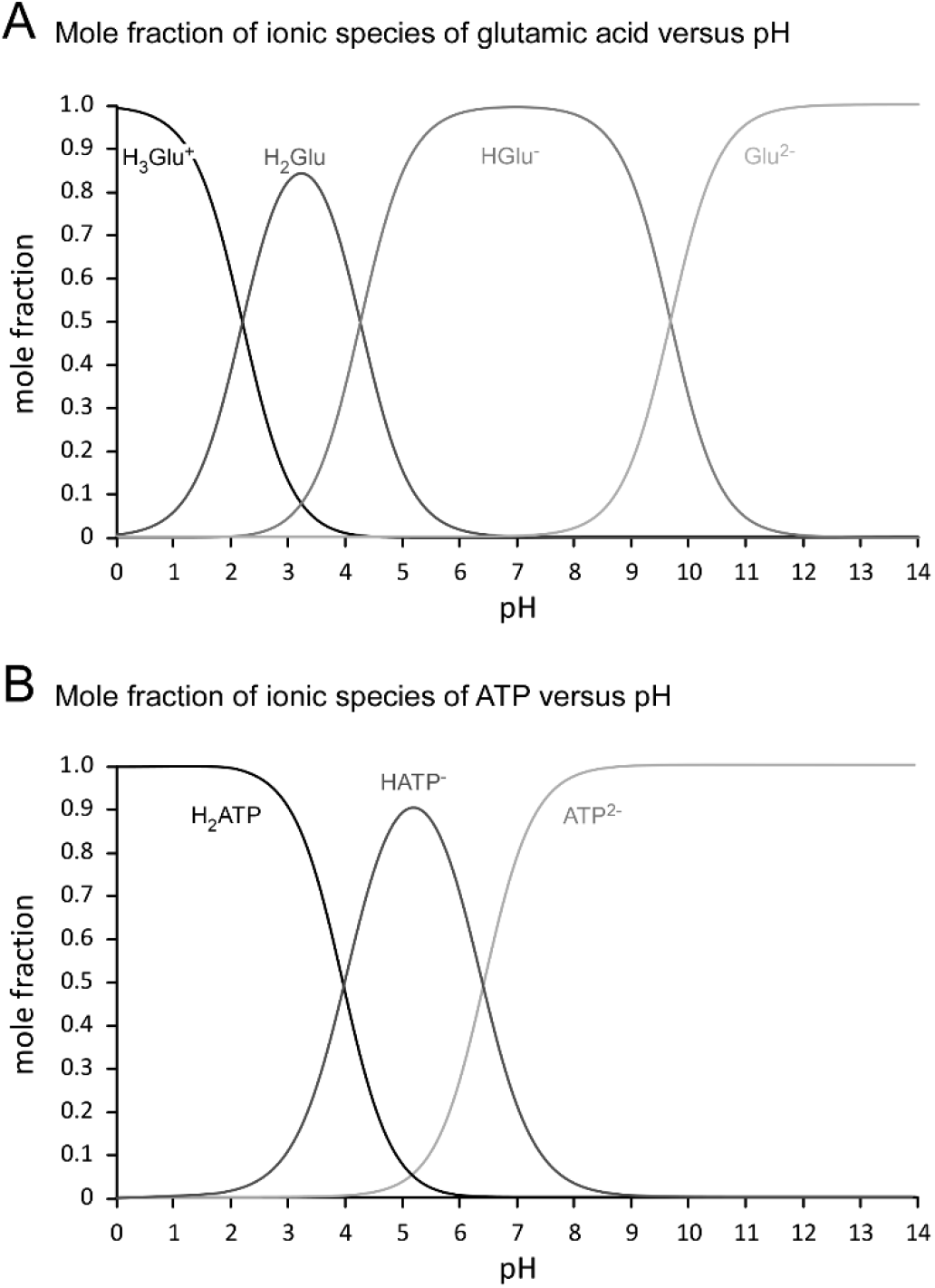
Bjerrum plots for glutamate and ATP. (**A**) Glutamate is a polyprotic acid, with three protonation sites, and at any pH the quantity of each ionizable species may be expressed as the ratio of the concentration of that species to the total concentration of glutamate. i.e. the mole fraction. The number of H^+^ that dissociate from glutamate after exocytosis can be calculated from the change in number of each species, and thus the change in net charge. (**B**) ATP has two protonation sites, and the same approach can be taken to calculate how many H^+^ dissociate from ATP after exocytosis.

**Figure S3.**
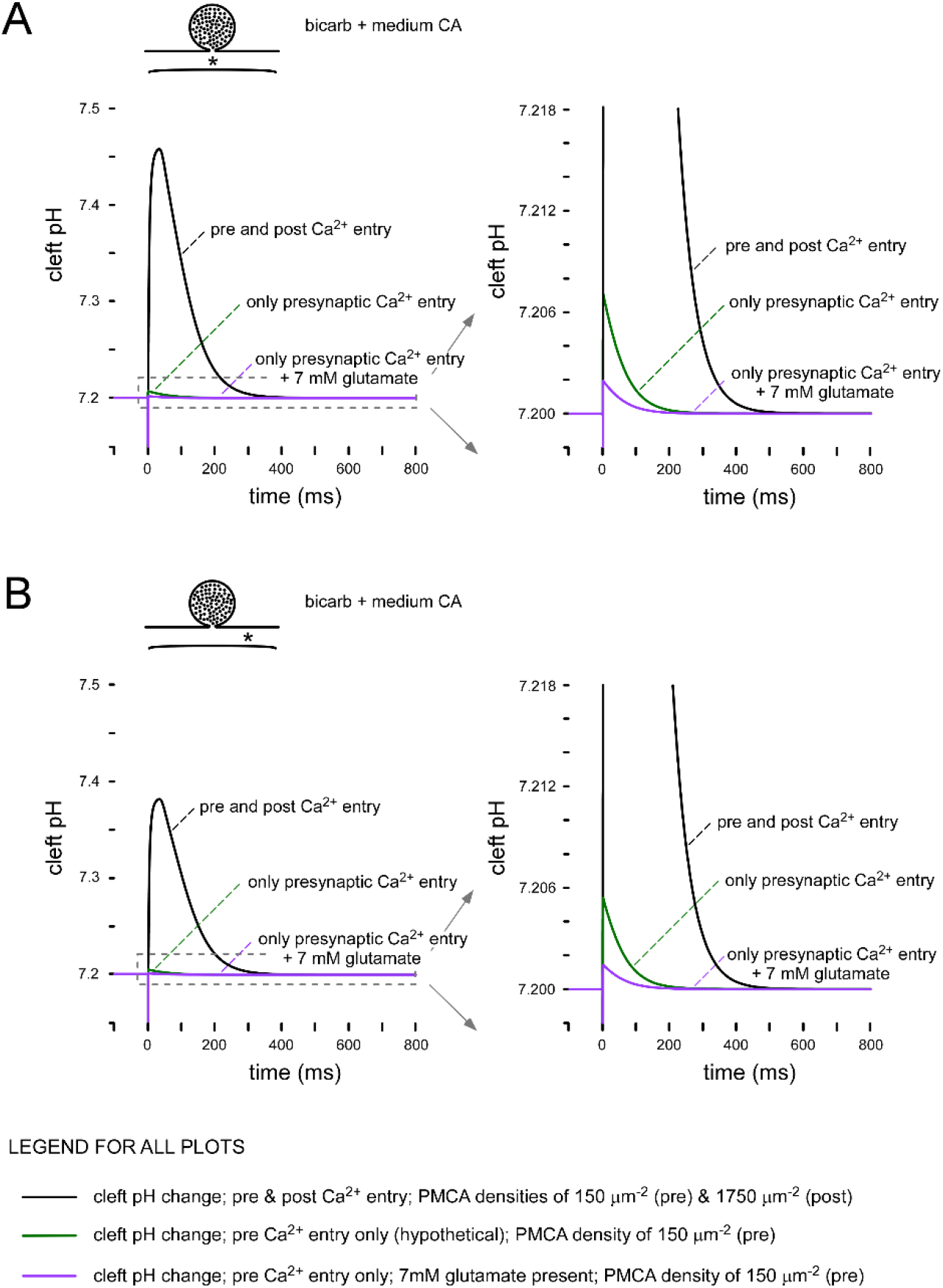
Substantiating presynaptic and postsynaptic PMCA density. (**A-B**) Output plots of the computational model demonstrating that the PMCA density values chosen for the pre- and postsynaptic membranes (n_pre_ and n_post_, respectively) yield pH transients that are compatible with empirical data published by Stawarski et al., 2020. In the presence of 7mM glutamic acid, a PMCA density of 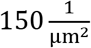 on the presynaptic side replicated the empirical data, i.e. an alkaline transient of only 0.0019 log units at the site of release (**A**), and 0.0014 midway along the cleft (**B**). In the presence of 7mM glutamate, a PMCA density of 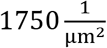 on the postsynaptic side replicated the empirical data, i.e. an alkaline transient of 0.258 log units at the site of release (**A**), and 0.181 midway along the cleft (**B**). The legend at the bottom applies to all plots. 19 mM bicarbonate and a medium level of CA activity is present under all conditions (600 En_acc_ nominal units). Note; the purple plot is hypothetical as we are unable to allow physiological levels of Ca^2+^ enter the presynaptic terminal without neurotransmitter release being triggered and a subsequent postsynaptic contribution to the pH transient (if 7mM glutamate is not present in the saline).

**Figure S4.**
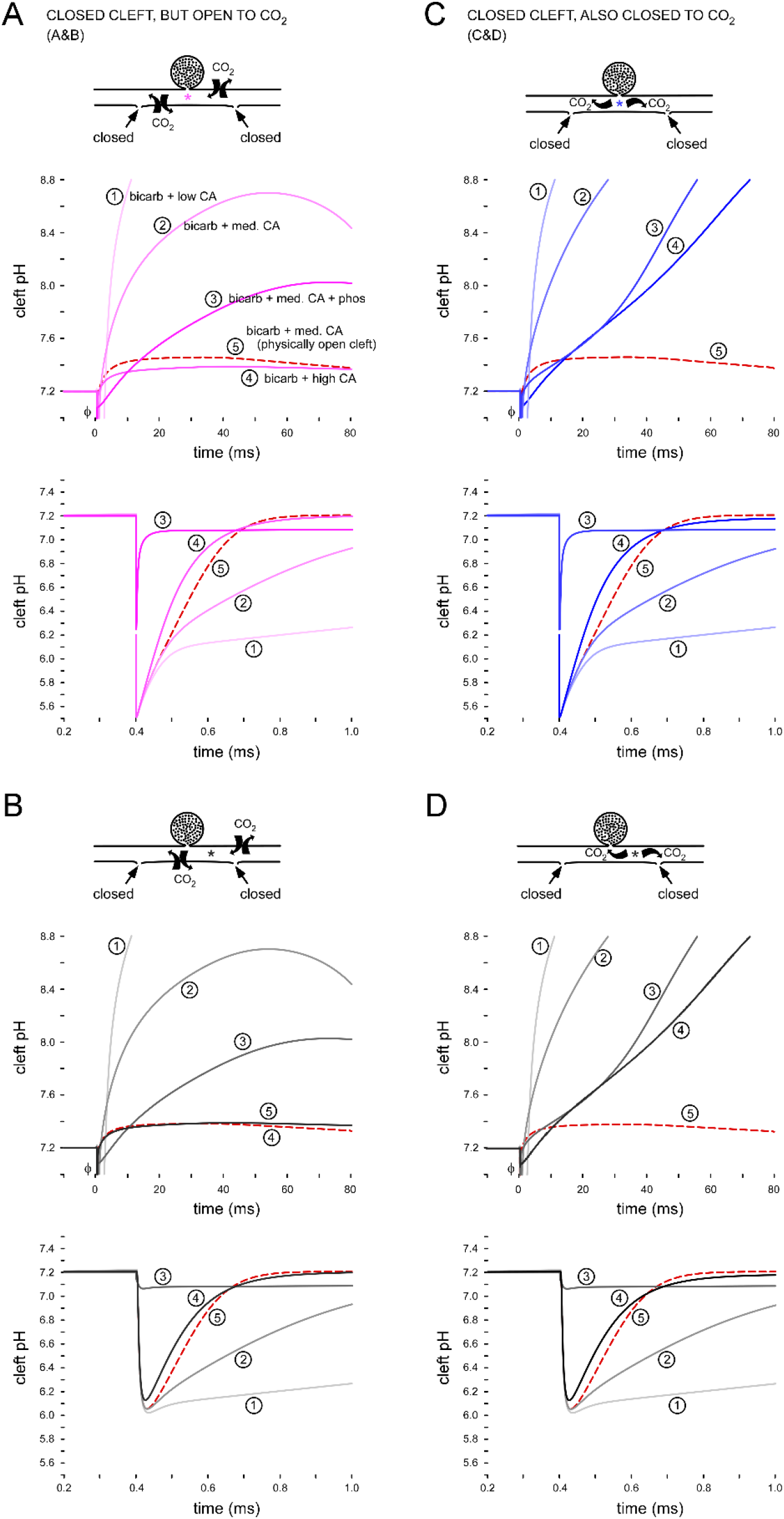
The influence of synapse morphology on cleft pH. (**A-B**) Detail of the output plots of the computational model showing the impact of preventing access from the synaptic cleft to the void of the SSR. Detail unavailable in **Figure 5**. The dashed line represents the control conditions, i.e. bicarbonate buffering in the presence of medium CA activity, full access to the SSR and membrane permeability to CO_2_. Low, medium and high CA activity are 100, 600 and 2,000 En_acc_ nominal units, respectively. pH 8.8 represents the pH at which the PMCA is expected to stall [33]. (**C-D**) Detail of plots showing the impact of both preventing cleft access to the SSR, and preventing CO_2_ movement across membranes. 8,000 glutamate molecules released and 19 mM bicarbonate buffering in all scenarios. Fusion pore opening (τ) is 1 μs. κ indicates truncated acidic transients.

## APPENDIX

To solve the diffusion equation numerically, we will discretize it in space to n concentric shells using a previous approach [31]. So h = R/n, r_j_ = jh, 0 ≤ j ≤ n,

We start with the equation

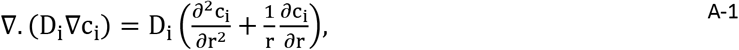

where we have assumed D_i_ is constant. Following [93], we can implement the finite difference approximation to discretize the diffusion equation as:
for (1 ≤ j ≤ n − 1)

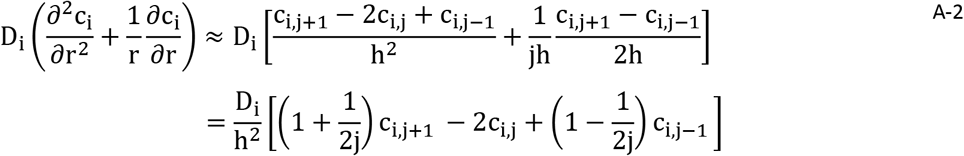

and for (j = 0)

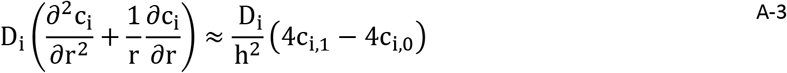

For simplicity let us define 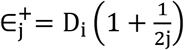 and 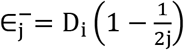 where 1 ≤ j ≤ n − 1.

To apply the boundary conditions numerically, we used a method described previously [73], which is to introduce virtual shells. For the outermost shell, we consider both possibilities of boundary conditions as discussed in the Methods. For the Neuman boundary condition (the case where there is no flux), we assume the virtual external shell is in equilibrium with the outermost shell, i.e.,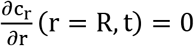, and, from there, we obtain 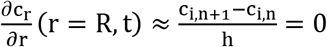 or c_i,n_ = c_i,n+1_. So, for (j = n) the Neuman boundary condition can be written as:

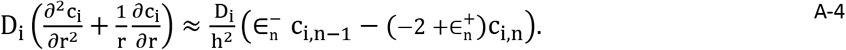

For the Dirichlet boundary condition (constant concentration field), we assume that in the external virtual shell, the concentration of all the species is constant during the simulation, i.e.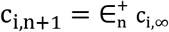. As a result, for (j = n) the Dirichlet boundary condition can be written as

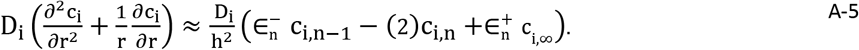

